# Bumblebee visual allometry results in locally improved resolution and globally improved sensitivity

**DOI:** 10.1101/380527

**Authors:** Gavin J. Taylor, Pierre Tichit, Marie D. Schmidt, Andrew J. Bodey, Christoph Rau, Emily Baird

## Abstract

The quality of visual information that is available to an animal is limited by the size of its eyes. Differences in eye size can be observed even between closely related individuals but we understand little about how this affects visual quality. Insects are good models for exploring the effects of size on visual systems because many species exhibit size polymorphism, which modifies both the size and shape of their eyes. Previous work in this area has been limited, however, due to the challenge of determining the 3D structure of eyes. To address this, we have developed a novel method based on x-ray tomography to measure the 3D structure of insect eyes and calculate their visual capabilities. We investigated visual allometry in the bumblebee *Bombus terrestris* and found that size affects specific aspects of visual quality including binocular overlap, optical sensitivity across the field of view, and visual resolution in the dorsofrontal visual field. This holistic study on eye allometry reveals that differential scaling between different eye areas provides substantial flexibility for larger bumblebees to have improved visual capabilities.

## Introduction

What an animal can see within its environment is restricted by its visual field, that is, the total angular region of the world from which light can be absorbed by the eye’s photoreceptors. To detect specific objects within this visual field, an eye needs spatial resolution, which is achieved through (and limited by) the arrangement of individual receptors that sample the spatial distribution of light (Land and Nilsson 2012). Having an eye with spatial resolution allows an animal to detect differences in light intensity reaching it from different directions, and the information this provides is crucial for the myriad of visually guided behaviours exhibited by different species (Cronin et al. 2014).

Over a given finite area, an eye cannot simultaneously maximise both its resolution and sensitivity (Land 1997) – to increase resolution, light must be sampled from a smaller region of space which necessarily reduces the amount of light captured, decreasing sensitivity (Snyder 1979). As a result, the relative density and optical properties of receptors often vary topologically across an eye, creating variations in visual resolution and optical sensitivity that ‘fine-tune’ the capabilities in certain regions of the visual field. We are familiar with this from our own vision – our fovea provides high-resolution over a small region of our visual field while our peripheral vision is blurred but covers a much larger region of the world (Land and Nilsson 2012). The eyes of other species have also evolved specializations to view regions of the world that have particular importance such as elongated regions of acute vision for detecting the horizon (Dahmen 1991) and ‘bright zones’ of high optical sensitivity for discriminating passing prey or potential mates against a bright background (Straw et al. 2006). Specialized areas thus represent a local investment in improving a specific visual capability that is related to an animal’s behavioural and ecological requirements. Although often overlooked, the periphery around specialized eye areas also provides important visual information that can, for instance, be used for wide-field motion detection (Franz and Krapp 2000). Thus, determining the topology of visual capabilities across an eye’s entire field of view (FOV) can provide important insights into the visually guided behaviours and environment of its owner (Moore et al. 2017) – for example, the topology of facet size (either with or without a region of enlarged facets) on the eyes of male bumblebees appears to indicate their species preferred mating strategy (perching or patrolling respectively) (Streinzer and Spaethe 2014).

Eyes are energetically costly to both develop and maintain (Niven and Laughlin 2008), so larger eyes represent an increased investment in vision. Across a wide range of animal groups, bigger individuals generally have absolutely larger eyes, although eyes do not typically grow linearly with body size but rather become proportionally smaller, even within a species (Jander and Jander 2002, Howland et al. 2004, Perl and Niven 2016). Increasing eye size should allow visual quality to improve – in a bigger eye, resolution can be increased by placing more receptors, or optical sensitivity can be increased enlarging the receptor size. Depending on the type of eye an animal has, achieving a given improvement in one parameter while maintaining the other requires a different relative size increase. A camera eye (a single lens focusing light onto many receptors) can increase its lens size in linear proportion to improved resolution to avoid losing sensitivity, while an apposition compound eye (where individual lenses focus light onto small groups of receptors) must grow in proportion to the resolution improvement squared, as it needs to both increase the size and number of its lenses (Land and Nilsson 2012). The relationship between the growth of any trait and an animal’s total body size is usually modelled using a power function^1^, with anatomical features that increase in size at a slower rate than the body being represented with an exponent less than 1. The camera type eyes of vertebrates on average scale with their body size to the power of 0.6 (Howland et al. 2004). Higher rates of eye allometry could be expected for invertebrate compound eyes given the limited returns provided by an increase in size but little is known about how these scale between taxa.

Scaling exponents for eye size within specific vertebrate groups do indeed exceed those of vertebrates and even include an unusual positive allometry rate in stingless Meliponine bees (Streinzer et al. 2016). Allometry studies within hymenopteran species – where body size can vary substantially between conspecifics – have shown that, while larger individuals of several ant species primarily invest in increasing their total facet number (Klotz et al. 1992, Zollikofer et al. 1995, Schwarz et al. 2011), other ants (Baker and Ma 2006, Perl and Niven 2016), and bumblebees (Kapustjanskij et al. 2007) increase both the number of facets and their size. Allometry has even been shown to vary across wood ant eyes, as differential scaling exponents of facet size are found between eye areas (Perl and Niven 2016) – leading to increased differences in the visual capabilities between specialized and peripheral eye areas. Given that homogenously increasing the size of a compound eye provides a relatively low improvement on its visual capabilities, we *hypothesise* that compound eyes are likely to scale differentially, such that the majority of the surface area of a larger eye will be invested to improve its vision in a small portion of its FOV.

To test if increasing compound eye size does indeed lead to the development or improvement of specialised visual regions in larger individuals, it is necessary to link the allometry of eye properties to the allometry of visual capabilities. Visual resolution in compound eyes is often estimated by dividing an assumed hemispherical FOV (Land 1997) by the number of facets, leading directly to the conclusion that resolution has the same (although negative) scaling exponent as facet number (Jander and Jander 2002). However, this assumption is not always supported by direct measurements of inter-ommatidial (IO) angle: IO angles from both desert ant and fruit fly eyes were found to have a lower absolute scaling exponents (−0.40 and −0.21 respectively) than the exponents for number of facets (0.75 and 0.58) (Zollikofer et al. 1995, Currea et al. 2018), while the scaling exponent of resolution was found to vary between different eye areas of Orange Sulphur butterfly eyes while the exponent of facet diameter remained constant (Merry et al. 2006). Furthermore, we calculated^2^ that a ~20% improvement in resolution of the mediofrontal area of bumblebee eyes is obtained given a 10% increase in total facet number (Spaethe and Chittka 2003). While these studies suggest that differential eye scaling leads to visual specializations, these analyses were made on specific compound eyes locations and thus do not reveal the full topology of their visual field.

Indeed, a major complication when comparing the corneal topology between different compound eyes is that eye shape can vary substantially, both between groups – for example, butterflies have nearly hemispherical eyes (Rutowski 2000), while flatter oval eyes are common in hymenopterans (Jander and Jander 2002) – and within groups – such as in the drastically different eye shapes found between male and female honeybees (Streinzer et al. 2013). In the absence of a common reference frame, topology based on general descriptions of eye shapes (e.g. the well-known dorsal rim area, (Labhart and Meyer 1999)) are not necessarily suitable for comparing visual capabilities between or within species, as corresponding anatomical areas may ultimately have different fields of view. Linking points on the eye to their projection into the visual world, and defining the visual capabilities they provide, is crucial for understanding how an eye samples information from the environment and how visually-guided behaviours, such as foraging, predation, and mating, are controlled. For example, the differences in the compound eye optics between different species of carpenter bees directly limits their optical sensitivity and thus the light levels at which they are able to forage (Somanathan et al. 2008, Somanathan et al. 2009). Similarly, bigger bumblebees have larger frontal facets enabling them to fly under dimmer illumination (Kapustjanskij et al. 2007) and to identify finer visual features (Spaethe and Chittka 2003) than smaller conspecifics. Whether these increased visual capabilities result solely from improvements to the frontal eye area, or if peripheral regions of the eye are also improved, remains unclear.

Here, we begin to explore the effect of size on visual capacity by comparing the topology across the entire eye of the size polymorphic bumblebee *Bombus terrestrìs*. The size differences of this species – which can equal an order of magnitude in mass (Goulson 2003) –influences some visually-guided behaviours (Spaethe and Weidenmüller 2002, Spaethe and Chittka 2003). Yet, little is known about how sensitivity and resolution across the entire visual field are affected. To test our hypothesis *that the differential scaling of compound eyes will primarily improve the visual capabilities in only a small region of the visual field*, we developed a novel method based on constructing 3D models of apposition compound eyes imaged with x-ray micro computed-tomography (microCT) (Baird and Taylor 2017). This allowed us to determine the visual field of an eye from the projection of its cornea into the world, and to then map the corneal facet topology across the FOV. Using this method, we could directly calculate the corneal IO angle topology on the world, and also the projected topologies of eye properties affecting optical sensitivity – namely, the facet diameter and the retinal and lens thicknesses. Crucially, this technique allows us to place eye properties and visual capabilities from different eyes (both intra- and inter-specific) into a common visual coordinate frame, enabling us to predict scaling exponents spatially across the visual field. We utilized our technique to investigate how size polymorphism influences *B. terrestris* vision by calculating the topology of eye properties and visual capabilities of six workers varying in linear body size by ×2.8 and in eye volume (EV) by ×3.9. As a reference, we also analyse eyes of European honeybees (*Apis mellifera*) that have a highly consistent worker size. Unlike *B. terrestris*, honeybee eyes have been the subject of extensive behavioural, anatomical, and physiological analyses, providing important data against which the results from our method can be critically compared. Overall, we find that our technique provides values that are consistent with those derived from other standard methods, indicating that it is suitable for comparing the visual capabilities between compound eyes. Moreover, we identify key features in the allometry of the visual topology of bumblebees that may represent general rules for how compound eyes scale with size.

## Methods

### Study animals

Bumblebees (*Bombus terrestris*) of small, medium, and large sizes, were obtained from a commercial hive (Koppert, UK), and honeybees (*Apis mellifera*) were collected from hives maintained at the Department of Biology, Lund University, Sweden. Several individuals of each species and size category were collected and anesthetized with carbon dioxide gas before dissecting.

### Sample preparation

Dissections were performed by removing either a bee’s left compound eye (to preserve this alone), or the front, bottom, and rear of its head capsule (to preserve the whole head). The inter-tegula width (ITW) was measured with digital callipers after dissecting the head as an indication of body size. The samples were immediately fixed in 3% paraformaldehyde, 2% glutaraldehyde, and 2% glucose in phosphate buffer (pH ~7.3, 0.2M) for 1 to 3 hours, and then washed in buffer before being immersed in 2% O_s_O_4_ for 1 hour to enhance the x-ray absorption contrast of soft tissues (Ribi et al. 2008). After washing the samples in buffer again, they were dehydrated with a graded alcohol series, and acetone was used to transition the samples to epoxy resin (Agar 100), which was cured in an oven at 60 ^o^C for ~48 hours. The samples in wet epoxy were placed on Perspex sticks, after which the external resin was peeled to expose the external cuticle of the sample (Taylor et al. 2016).

### X-Ray Microtomography

Tomographic imaging was conducted using synchrotron light at the Diamond-Manchester Imaging Branchline I13-2 (Rau et al. 2011, Pešić et al. 2013) at the Diamond Light Source, United Kingdom. At the beamline, an undulator (gap set to 5 mm) was used to produce a polychromatic (5 to 35 keV) ‘pink’ beam of partially-coherent, near-parallel, x-rays with lower energies suppressed using metal filters. We collected projection images of each sample from 4001 equally spaced angles over 180° of continuous rotation with a scintillator coupled pco.edge 5.5 (PCO AG) detector. A propagation (sample to scintillator) distance of 100 mm was used to give a moderate level of inline phase contrast. Dissected heads were imaged using ×2.5 total magnification (2.6 *μ*m effective pixel size, Fig. 1A), while isolated eyes were imaged with ×4 total magnification (1.6 *μ*m effective pixel size, Fig. 1B,C). Projection images were flat- and dark-field corrected prior to reconstruction into 3D volumes using a filtered back projection algorithm (Basham et al. 2015, Titarenko 2016).

**Figure 1:**
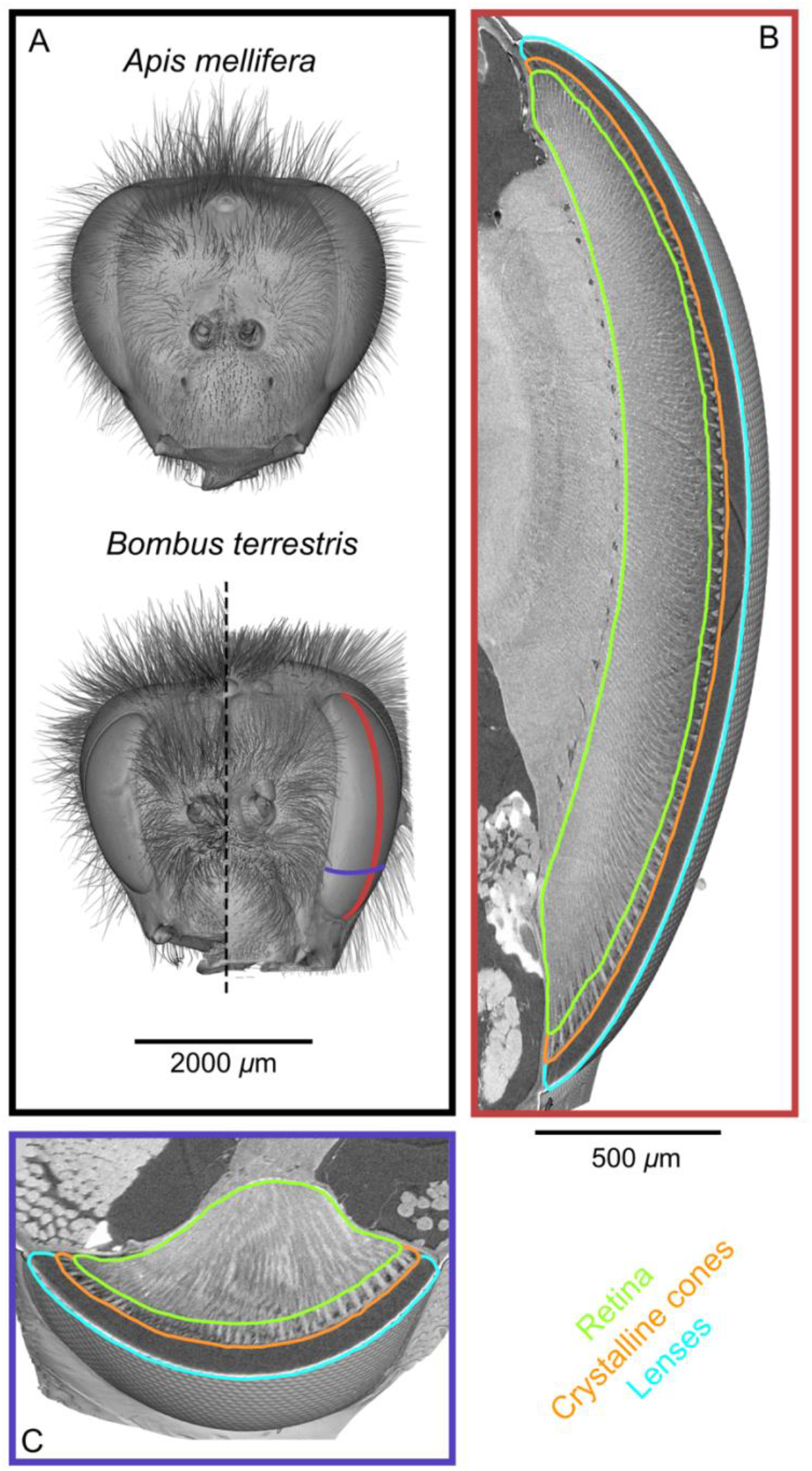
Tomographic images of bee heads and eyes. **A**, volume rendering of the intact heads of workers from the bee species used in this study. The right-hand side of a small *B. terrestris* head is shown (ITW 2.94 mm) in comparison to left hand side of a large individual from the same colony (ITW 5.42 mm, note that the mandibles and some hair of this bee were too large to image). The ITW of the *A. Mellifera* specimen was 3.55 mm. **B**, vertical section along the mid-line of a *B. terrestrìs* apposition compound eye showing the gross morphology of the lenses, crystalline cones, and retina. A portion of the optic lobe is also visible to the left of the retina. **C**, a transverse section across the ventral portion of the compound eye showing the same features as in B. The approximate location of the sections in B and C are indicated with lines on the larger *B. terrestris* eye in A; both B and C have the same scaling.

### Volumetric analysis

The reconstructed 32-bit volumes were initially cropped spatially (around the scanned feature), and in their intensity range using the program Drishti Paint (Limaye 2012), and resaved as 8-bit files. Amira (FEI) was used for further analysis of the volumes in three ways; i) by manually labelling the structures of a compound eye (Fig. 1B,C), ii) alignment of the labelled compound eyes of a given species onto the scan of a full head (Fig. 2A), and iii) measurement of facet dimensions on a compound eye (Fig. 2B). Volumes for labelling and registration were typically resampled to 5 *μ*m voxel size using a Mitchel filter, however, resampling was not used for facet measurement as this could prevent the facets borders from being visualized. Additional details about the procedure used to process volumes in Amira are provided in the Supplemental Methods.

**Figure 2:**
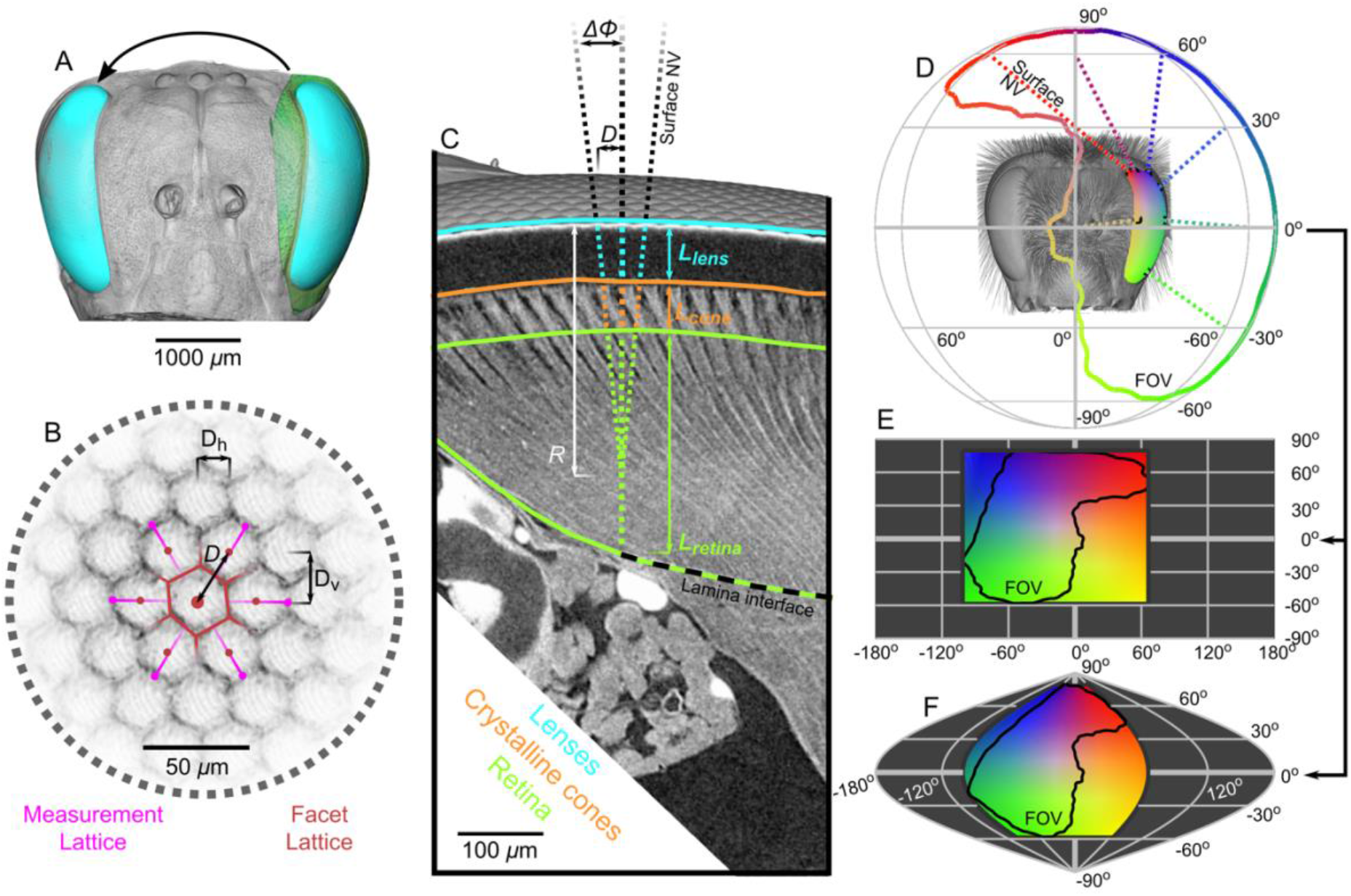
Visual analysis of performed on tomographic scans of bee eyes. **A**, a volume rendering of the left compound eye (green) of each bee was aligned onto a rendering of the full head of another bee (grey), which allowed the segmented eye (cyan) to be placed relative to the head and also mirrored to the right side. **B**, to measure the local facet dimensions, six points were selected on the opposing borders of the six facets that surrounded a central facet. These points were then used to compute the local structure of the facet lattice from which the lens diameter (*D*, the horizontal *D_h_* and vertical *D_v_* facet dimensions were not used in this study) could be calculated. **C**, surface NVs were calculated from the exterior surface of the compounded eye lenses to indicate the local viewing direction and the difference between viewing directions at a distance of one lens diameter provides the corneal IO angle (*ΔΦ*, the corneal radius of curvature *R* was calculated as *D*/*ΔΦ*). The NV was traced into the eye and the points where it intersected the front surface of the CC, the retina, and the lamina interface were used to determine the thickness of these structures (*L_lens_, L_cone_*, and *L_retina_* respectively). Observe that the angle of the CC and the photoreceptors in the retina can be misaligned from the corneal NV. **D**, the projection of the NV_s_ (several are plotted as dotted lines) from the eye onto a sphere indicates the extent of the eye’s FOV, which can also be represented on equirectangular (**E**) or sinusoidal (**F**) projections on which the FOV is indicated by black lines. The colour coding in D, E, and F indicates the viewing direction, and was applied by stretching a 2D colourmap across the FOV extent on the equirectangular projection in D. Thus, colours on both the surface NVs, eye, and FOV in D indicate equivalent viewing directions for matching colours on the projections in E and F.

### Computational analysis

We developed Matlab scripts to compute the measurements reported in this study from the volumetric analysis performed in Amira on apposition compound eyes. This procedure used the labelled volumes, facet measurements and transforms to compute the eye properties: eye surface area, eye volume, facet number, *facet diameter, radius of curvature*, and *thicknesses* (for the lenses, CC, and retina); visual capabilities: corneal FOV, binocular FOV, complete FOV, *corneal IO angle*, and *optical sensitivity*; and the derived metrics: *eye parameter*. The italicized variables were determined locally, that is, they were calculated at sampling points that were equally spaced at 25 *μ*m intervals across each bee’s corneal surface. The corneal normal vector (NV) of each sampling point was determined as an indication of the viewing direction of that part of the eye in space (Fig. 2C). NVs were calculated by fitting a 2^nd^ order 2D polynomial directly to the local voxels of the corneal surface around each sampling point (Taylor et al. 2016). While sampling points were equally spaced on a given eye, their projected NVs did not necessarily have equal angular spacing because the eyes radius of curvature varied. To ensure uniform angular sampling of each variable, we generated a single set of world points that were equally distributed onto a sphere (spaced at ~1° intervals). For each bee, we determined which world points were inside its FOV (Fig. 2D), before using the viewing direction of the sampling points to interpolate each locally calculated variable onto the world points. This allowed locally calculated variables to be represented in both eye- and world-centric coordinates, both of which could be used to represent spatial variation in a variable using a topology. However, only the world represents a common reference frame to allow direct comparisons for the topology of a given variable between multiple individuals and species (Fig. 2E,F). Additional details about this computational analysis procedure and a discussion on its limitations are provided as Supplemental Methods and the original Matlab scripts are available on request.

### Allometry

Allometry is described by fitting the parameters *b* and *a* in the power function *Y*=*bx^a^* (Huxley and Teissier 1936) after logarithmic transformation of the size indicator (*x*) and the dependent variable (*Y*). The transformation results in the equation log_10_(*Y*) = log_10_(*b*) + *a*log_10_(*X*), for which the parameters can be obtained from the least-squares fit of a 1^st^ order polynomial function. All variables are converted to linear measurements before the logarithmic transform (by taking the square root of areas and the cube root of volumes). Calculating the linear correlation of the transformed variables also provides R^2^ and *p* to indicate the effectiveness of a linear fit at describing the data. Functions with non-significant (*p* > 0.05) correlations are plotted as dotted lines. We usually used eye size as the independent parameter when measuring visual allometry as we wished to focus on investigating how, given an initial total investment in eye size, the resolution or sensitivity of an eye was improved. In other contexts, it may be desirable to use body size as the independent variable, which can be accomplished by multiplying the scaling exponent measured for a specific variable vs. eye size by the scaling exponent for bumblebee eye size vs. body size (0.45). We calculated allometry functions for the facet-wise mean values of variables for bumblebees of different sizes (Table S1). We also calculated functions locally for variables based on their projection into the world from each eye and represent these as spatial maps of the scaling exponent of each variable (Figs. 7 & S5). Functions were only fitted for points in space viewed by at least four individuals, which totalled 2473 of the 10242 world points. The topology of all variables in our analysis show some level of spatial autocorrelation and indicating that fewer parameters could be used to describe the observed variation.

## Results

Bumblebee eyes display negative allometry, increasing in both area and volume at a slower rate than their body size (Fig. 3A, Table S1). Their corneal FOV – the angular projection of the cornea onto the world – also increases with eye size (Fig. 3C), although at a slower rate than the total number of facets (Fig. 3B). Naively, this suggests that bigger bumblebees would have more facets per unit area, implying lower corneal IO angles. However, our calculations suggest that corneal IO angles generally maintain a relatively similar, although broad, distribution as eye size increases, with only a small decrease in the mean IO angle present in the largest bees. The lowest IO angles were slightly less than 1° for all bees (Fig. 3D). Our facet-based average for IO angles are somewhat lower than the values generated by the hemispherical assumption (triangles, Fig. 3D), but are higher than would be predicted by dividing the calculated corneal FOV by the number of facets (stars, Fig. 3D). Nonetheless, the results of both assumptions do lie within the range of observed values for each bee (grey histograms, Fig. 3D). The IO angle (*ΔΦ*) is itself derived from local eye properties: the facet diameter divided (D) by the radius of curvature (R). When considering the allometry of these eye properties, we found that both facet diameter (Fig. 3E) and radius (Fig. S1A) clearly increase with eye size. As both properties had a similar scaling exponent (Table S1), this results in similar IO angles across the range of *Bombus* eye sizes examined.

**Figure 3:**
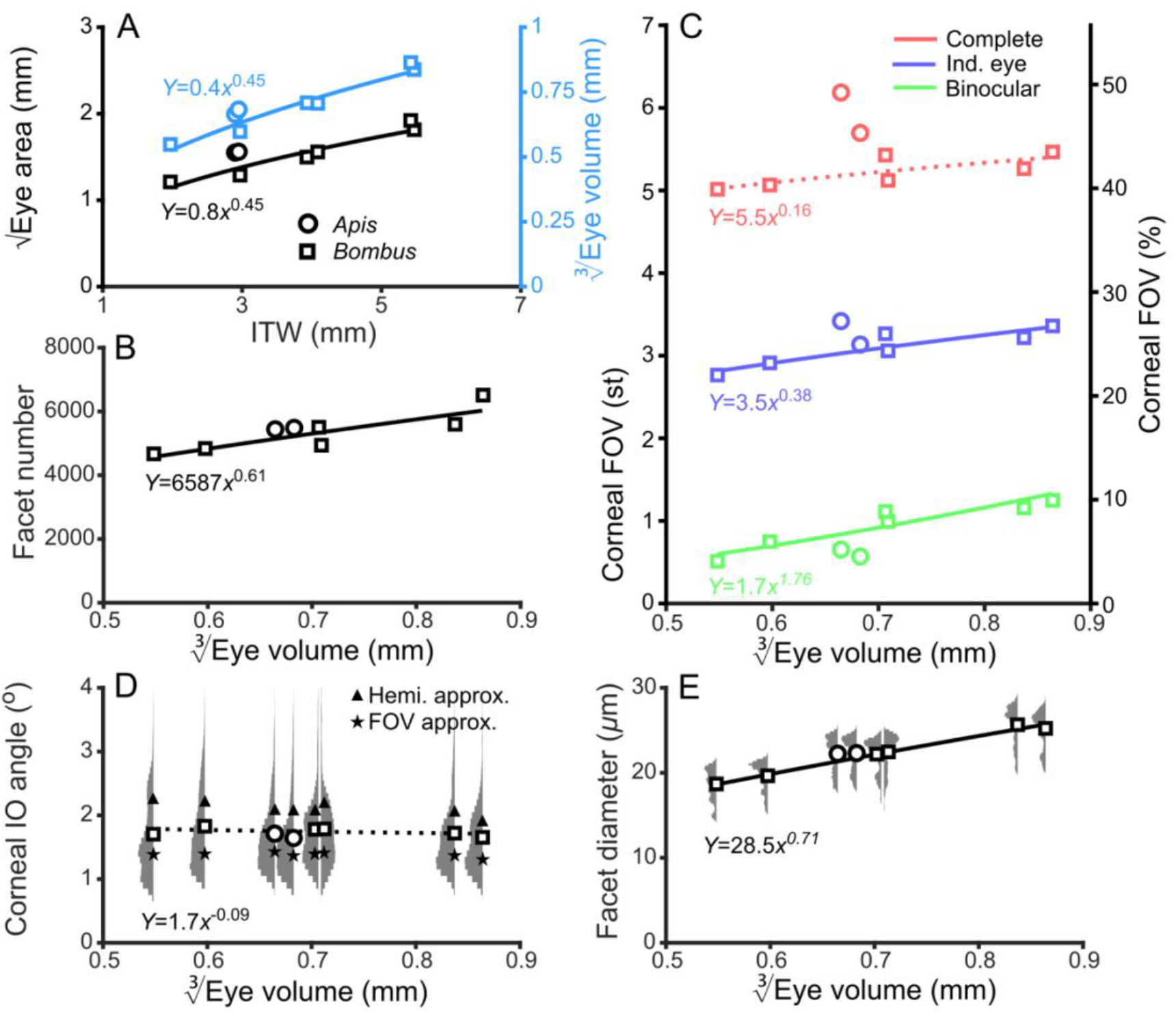
Standard variables for describing compound eyes. **A**, √Eye area and 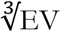 (on the right-hand Y-axis) vs. ITW. **B**, the number of facets in each compound eye as a function of 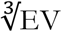. **C**, the size of the corneal individual, binocular, and complete FOVs both in steradians (left Y-axis, max: 4*π*) and as a percentage of the total visual sphere (right Y-axis, max: 100%). **D**, average corneal IO angles and relative distribution for each bee. The triangles indicate the result of dividing a hemisphere by the total facet number to predict average IO angle (Hemi. approx.) (Land 1997), while the stars indicate the results of dividing an individual eyes angular FOV by its total facet number (FOV approx.). **E**, averages and relative distributions as for D, but for measurements of the facet diameters for each bee. Squares denote bumblebees and circles denote honeybees in all plots, while colour coding is indicated on each panel. A power function was fitted to the *Bombus* measurements for each parameter (Table S1); the resulting functions are written and plotted in each panel. The dotted lines in C and D indicate that the correlations between those parameters and 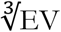 are not significant (Table S1).

### Fields of view

The corneal FOV increases with bumblebee eye size (Fig. 3C), but it appears this increase does not result from simply enlarging the FOV in all directions (Fig. 4A,B). The dorsolateral limit of the FOV of each eye is relatively consistent (between 30° to 60° elevation (el.) and −90° to −60° azimuth (az.), Fig. 4B), while bigger bees appear to enlarge their FOV dorsoanteriorly (between 0° to 90° el. and −15° to 75° az.) and to a lesser extent ventrolaterally (between 0° to −60° el. and −105° to −60° az.). When the FOV of each bee’s right eye is also considered, it is apparent that all bees have regions of corneal binocularity (Fig. 4C), that increase in angular area with eye size (Fig. 3C). While the increase in the binocular FOV is primarily observed dorsoanteriorly (between 30° to 90° el. and −75° to 75° az., Fig. 4C), a region of binocularity is also observed facing directly forwards (between −30° to 30° el. and −15° to 15° az.). Given their shapes, these appear to be two distinct binocular regions; they are separated on the smallest bee and merge as the binocular field is enlarged on bigger bees. The corneal FOV is a spatial, but binary, representation of each bee’s vision across which the eye properties and the resultant visual capabilities vary topologically.

**Figure 4:**
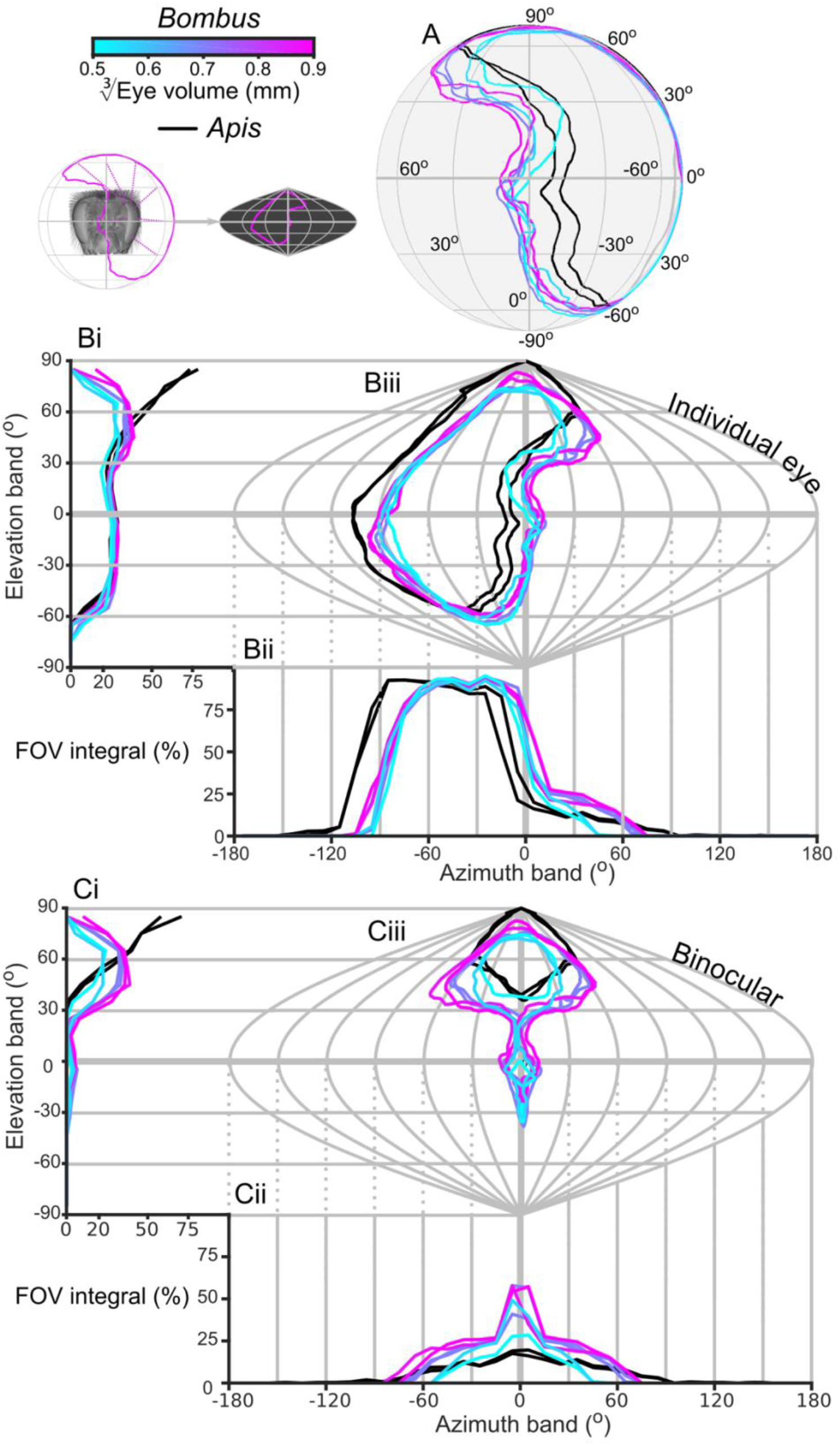
The corneal visual fields of compound eyes. **A**, the corneal FOV of each bee’s left eye shown on a sphere representing the world. **B**, a sinusoidal projection and analysis of the FOVs (iii). The integrated corneal FOV profiles are shown across elevation (i, the integral of all azimuth points in the FOV as a function of elevation) and azimuth (ii, the sum of all elevation points in the FOV as a function of azimuth) and expressed as a percentage of the total number of points. **C**, as for B, but depicting the limit of the corneal binocular overlap between the visual field of left and right eyes for each bee. The cyan-to-magenta colour bar indicates 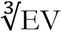 of *Bombus* for each curve in all panels, while black lines indicate honeybees. See Fig. 2D,F for an example of the relationship between a sphere and its sinusoidal projection.

### Optical sensitivity

Optical sensitivity is related to facet diameter and the thickness of the underlying retina (Warrant and Nilsson 1998) and both of these properties increase with eye size (Figs. 3E & 5A). This suggests that larger bees have more sensitive ommatidia than smaller bees, if the acceptance angles of receptors are assumed to be equivalent to the IO angles calculated (Fig. S5D). Taking into consideration the similar eye-wide averages for IO angle (Fig. 3D), an increase in retinal thickness suggests that bigger bees invest in improving their sensitivity rather than their visual resolution. However, there is substantial variation in the histograms of the measured variables for each eye (Figs. 2D,E & 5A), so it is of further interest to investigate the topology of how these are projected into the visual field.

**Figure 5:**
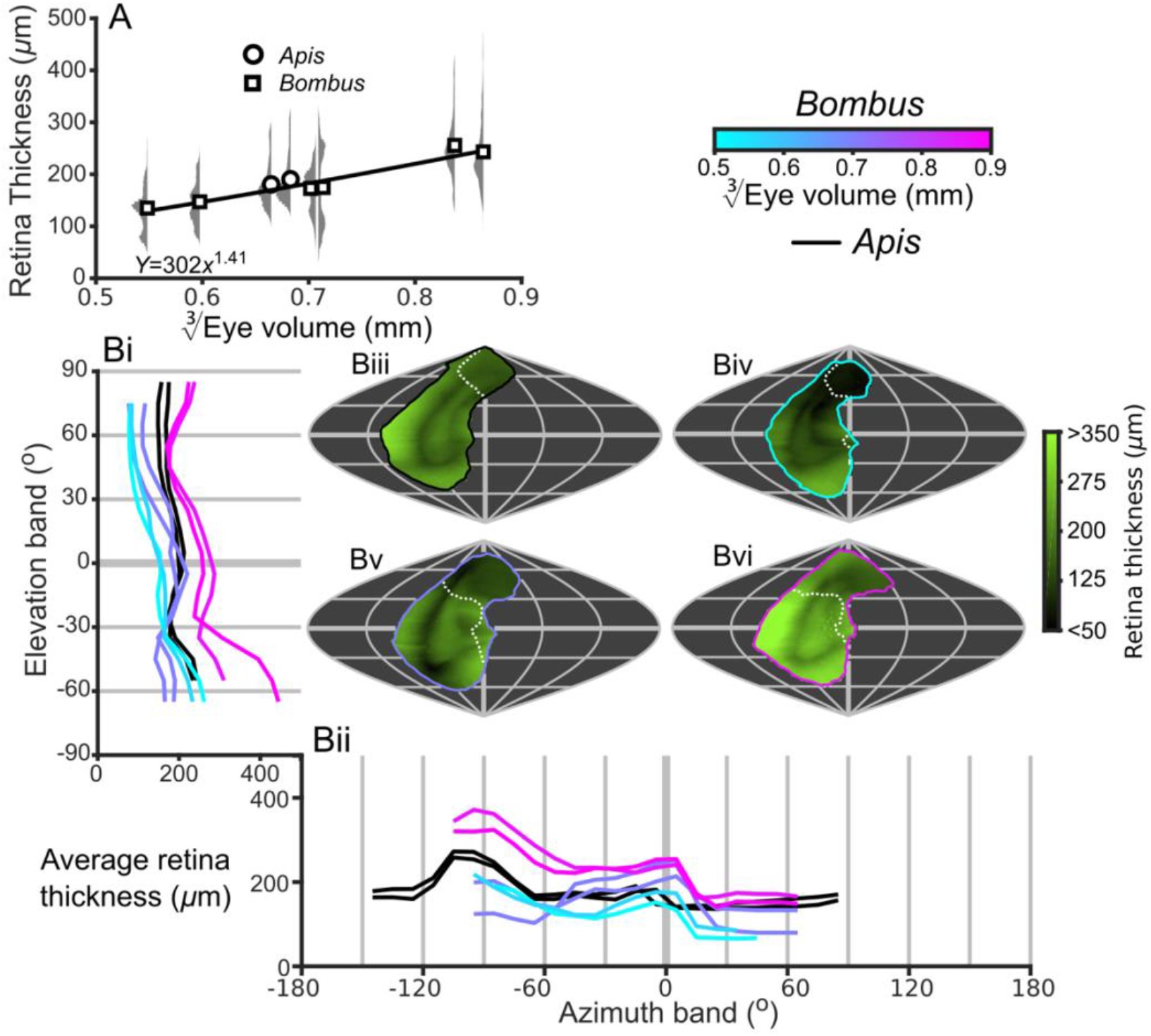
Description of the retinal thickness underlying compound eyes. **A**, the average retinal thicknesses and relative distribution for each bee as a function of 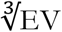 (notes in the caption for Fig. 3E also apply to this panel). **B**, the topographic distribution of retinal thickness (indicated by the green colour bar) projected from each bee into the visual world, shown as sinusoidal projections for a honeybee (iii), and small (iv), medium (v), and large (vi) sized bumblebees (azimuth and elevation lines are plotted at 60° and 30° intervals respectively). The average retinal thickness profiles are shown for elevation (i, thickness is averaged across *all* azimuth points in each eye’s FOV as function of elevation) and azimuth (ii, thickness is averaged across *all* elevation points in each eye’s FOV as a function of azimuth). The white dotted line across the topology in Biii to Bvi indicate the border of the bee’s corneal binocular FOV, and the same colour scheme is used as in Fig. 4 to indicate a bee’s EV and species.

### Projected topologies

The lowest corneal IO angles within each bee’s FOV are observed in a laterally positioned vertical band running from −45° until 60° el. at approximately −60° az., while the highest IO angles are observed at the posterior and rightmost dorsal limits of the visual field (Fig. 6A). All *Bombus* have a similar average IO angle profile across elevation (all average ~1.5° between −30° to 30° el., Fig. 6Ai). When averaged across azimuth, it is apparent that larger eyes have higher frontal resolution (between −45° to 15° az., Fig. 6Aii), although the lowest azimuthal average IO angle is *not* directed frontally, but rather at ~-60° for all bees. A distinctly different topology is present when projecting the facet diameters into each bee’s FOV (Fig. 6B). Larger *Bombus* have larger facets (Fig. 3E), each bee’s facet diameters (averaged across elevation) are highest ventrally (<0°, Fig. 6Bi) and lowest dorsally (>45°). While the diameters for each bee are generally similar within these elevation ranges, they are connected by a transitory range as diameters decrease from 0° to 45°. When averaging facet diameter across azimuth, each *Bombus* has its largest facets facing towards the lateral FOV (<60°, Fig. 6Bii). Retinal thickness is projected into visual space with a similar, although less consistent, topology as facet diameter; average thickness generally increases towards their ventral and lateral FOVs (Fig. 5B). Interestingly, lens thickness has a different topology to the other properties we have described, as it peaks in the frontal visual field (Fig. S2Bii) and is also correlated to facet diameter (r = 0.65, Fig. S6). After examining the projection of these variables into visual space, it appears that bumblebee eye properties maintain a similar topology that essentially scales with eye size (facet diameter (Fig. 6B), radius of curvature (Fig. S1B), retinal thickness (Fig. 5B), lens thickness (Fig. S2B), CC thickness (Fig. S3B)). However, the IO angle is a function of local facet diameter and radius of curvature, clear variations in the local topology of corneal IO angles (Fig. 6A) evidently arise from subtle changes in eye properties.

**Figure 6:**
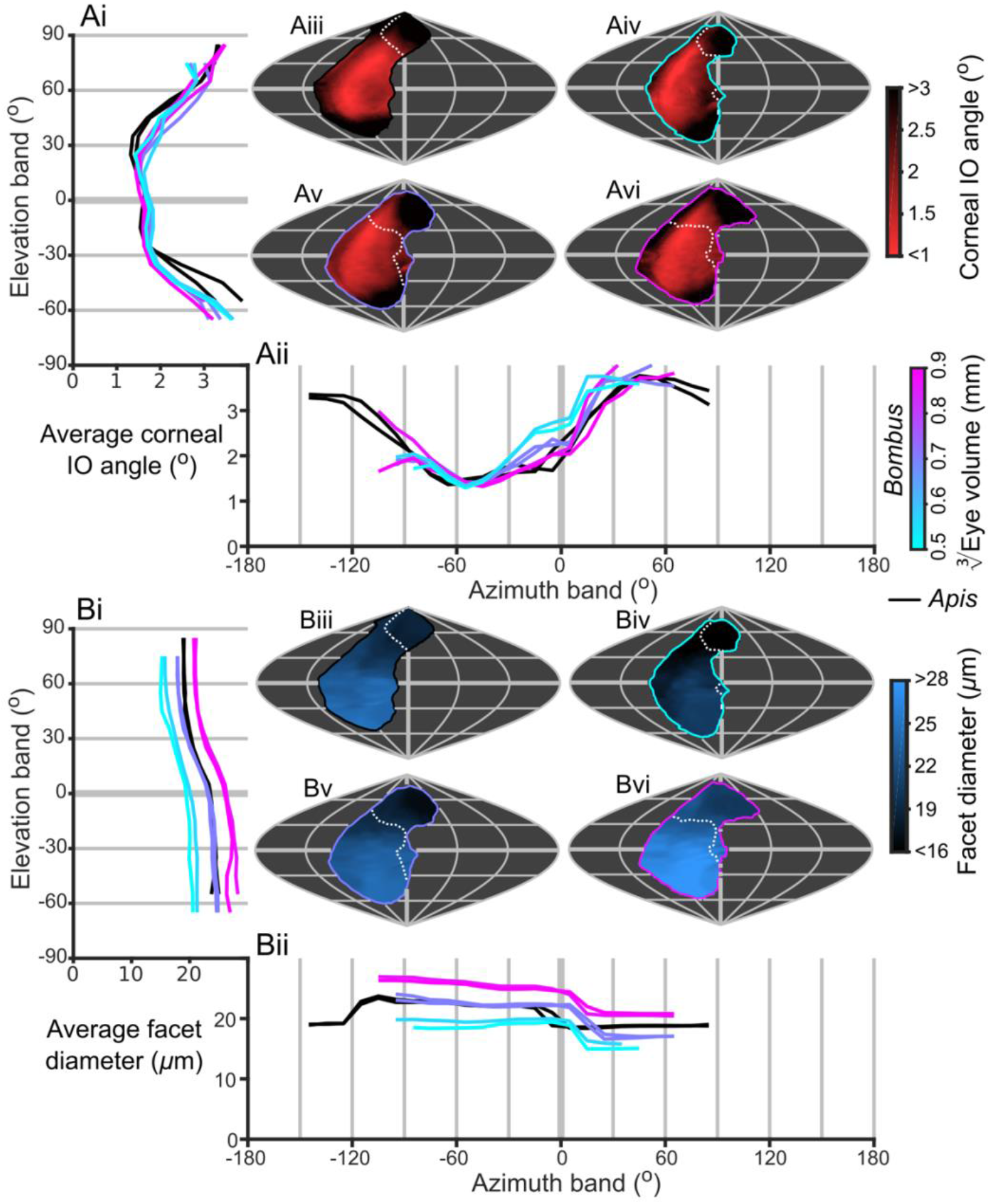
Projection of corneal IO angle and facet diameter into the visual world. **A**, the topographic distribution of IO angle (indicated by the red colour bar) projected from each bee’s eye, shown as sinusoidal projections for a honeybee (iii), and small (iv), medium (v) and large (vi) sized bumblebees (azimuth and elevation lines are plotted at 60° and 30° intervals respectively). The average IO angle profiles are shown for elevation (i, where IO angle is averaged a function of elevation, as described in the caption for Fig. 5B) and azimuth (ii, where IO angle is averaged as a function of elevation). **B**, as for A, but showing the topology (indicated by the blue colour bar) and average profiles of facet diameter. The white dotted line across the topology in Biii to Bvi indicate the border of the bee’s corneal binocular FOV, and the same colour scheme is used as in Fig. 4 to indicate a bee’s EV and species.

### Mapping scaling rates

To specifically examine how changes in eye size affect the allometry of the topology of different characteristics, we calculated maps of the local scaling exponents, for the eye properties and visual capabilities in the common coordinate frame provided by the visual world. As identified from the corneal IO topology (Fig. 6A), the scaling exponent maps show that larger bees improved their resolution in their frontal and dorsofrontal FOVs (between −15° to 60° el. and −30° to 10° az., Fig. 7A). However, as the facet diameter maintains an almost uniform exponent across the visual field (Fig. 7B), local variations in the scaling rate of the radius of curvature (Fig. S5C) cause differences in the IO exponent across the eye. In larger bumblebees, changes in the eye’s radius, and consequently its shape, are the primary determinant of changes in corneal IO angle. The local corneal IO angle has strong negative correlation to the radius of curvature (*r* = −0.84) but is not correlated to facet diameter (*r* = −0.07, Fig. S6). Both retinal thickness and facet diameter contribute to sensitivity, although the local scaling exponent of retinal thickness has substantially more variation (Fig. 7C); it is highest in the dorsal hemisphere and, surprisingly, lowest in the frontal region associated with binocularity (Fig. 4C). Subjectively, these maps of scaling exponents provide a clear indication about where and how the topology of bumblebee eye properties changes as a function of eye size (Figs. 7 & S5), even more so than the actual topologies themselves (Figs. 4B, 5, & S1–4B).

**Figure 7:**
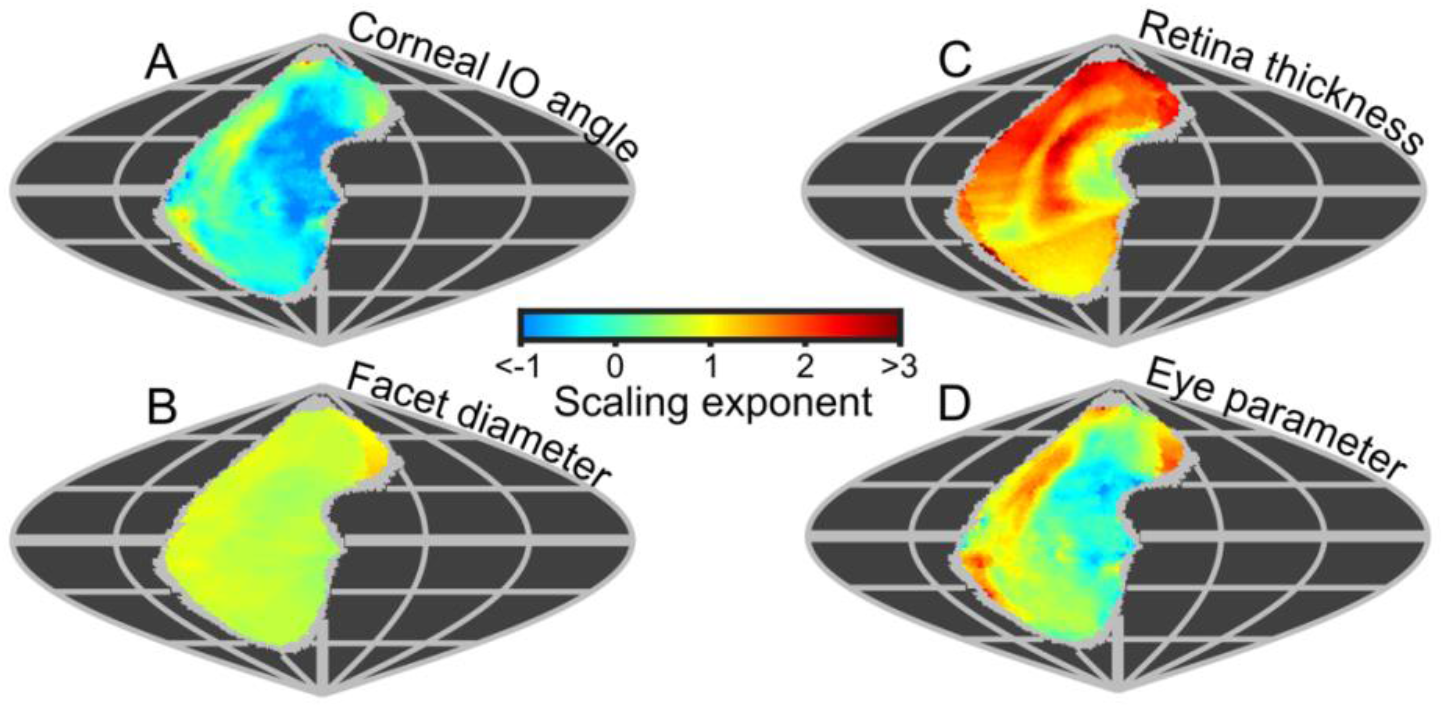
Eye metrics and maps of scaling exponents in the visual world. **A** to **E** show maps of the scaling exponents calculated for bumblebees for the variable specified on each map. The scaling exponents for all variables use the blue-to-red colour bar. Positive scaling exponents (red) indicate visual ‘improvement’ when eye size increases for the facet diameter (B), retina thickness (C), while conversely, the corneal IO angle (A) improves with eye size given a negative exponent (blue). A positive exponent for eye parameter (D) indicates increasing optical sensitivity, while a negative exponent indicates improving optical resolution. We limited the scaling exponent calculations to regions viewed by at least four bees, and the grey fringe in each map indicates the additional regions (viewed by three or fewer individuals) that were not used in the calculations.

### Eye parameter

Until now, we have described eye properties and their resultant visual capabilities to investigate how bumblebees invest in vision as their eye size increases. The eye parameter (*P*) is an additional metric (calculated as the local IO angle multiplied by the facet diameter, (Snyder 1979)) that provides an indication about whether facets are optimized for resolution (where *P* approaches 0.29 if the resolution of the eye is limited by the diffraction of green light) or sensitivity (where *P* may be 1 or higher). Although the average eye parameter increases with eye size (Fig. S4A), we found all bumblebees had a similar average eye parameter profile between −30° and 10° az. (Fig. S4Bii). Scaling exponent maps showed that the eye parameter of larger bees was slightly reduced in a similar region to where their IO angles decreased, but increased in the periphery of the FOV (Fig. 7A,D). This indicates that the dorsofrontally improved IO angles correspond to a region of improved optical resolution on larger bees which strongly suggests that this region of the eye represents an acute zone optimized for high visual resolution (Land 1997).

### Honeybees

Honeybees have slightly larger eye areas and volumes relative to their ITW than bumblebees (Fig 2A). When compared on the basis of their EV, the average values for honeybee eyes are most similar to those of medium sized *Bombus* (in terms of corneal IO angle (Fig 2D), facet diameter (Fig 2E), radius of curvature (Fig. S1A), rhabdom length (Fig. 5A), and eye parameter (Fig. 7F)). Two distinguishing characteristics of honeybee eyes are that their values for corneal lens thickness are distinctly lower than all bumblebees (Fig. S2A), and the extent of their binocular overlap is also reduced compared to medium sized *Bombus* (Fig 2C; as the FOV of each individual eye is similar, this results in a larger complete FOV). The corneal visual fields of honeybees tend to be shifted dorsoposteriorly relative to those bumblebees (Fig. 4A,B), while their binocular overlap is limited to their dorsal visual field and does not extend anteriorly (Fig. 4C). After accounting for the differences in their visual fields, the projected topologies of the honeybees are most similar to those of medium sized bumblebees, with two differences. Firstly, honeybees have an obvious increase in retinal thickness in their lateral visual field (Fig. 5Bii, −120° to −75° az.) and, secondly, they have relatively low IO angles in their frontal visual field (Fig. 6Aii, −30° to 30° az.), which is the approximately the region where larger bumblebees increase their visual acuity.

## Discussion

In this study, we demonstrated a new microCT-based method for reconstructing the visual world of insects with unprecedented detail. We created 3D models directly from the volumetric images of bee’s apposition compound eyes to determine how their surfaces are projected into the visual world. This allowed us to determine each eye’s corneal field of view, across which we could calculate the topology of the both corneal IO angle and of properties influencing optical sensitivity, including lens size, and rhabdom and lens thickness. As our method enabled us to project the FOV from each eye into the world, we could map the scaling relationships of eye properties and visual capabilities for bumblebees. This enabled us to disambiguate how bumblebees of different sizes invest in resolution and sensitivity across their visual field, revealing unexpected patterns in how this investment changes with eye size.

### Comparison of eye analysis techniques

Anatomical studies of insect eyes have typically been based on measurements of 2D representations – either from imaging histological cross sections or from flattened replicas of the eye surface (Ribi et al. 1989). While these techniques do allow optical dimensions to be measured from the eye’s morphology, they lose the 3D location of these measurements relative to the remainder of the eye and head. Some studies on compound eyes also measure 2D approximations of 3D shapes, such as the projected surface area of compound eye from a given viewpoint (Spaethe and Chittka 2003, Kapustjanskij et al. 2007, Perl and Niven 2016). While a projected eye area is dependent on its actual surface area^3^, the former is not broadly optically relevant, and the two measurements are not always clearly distinguished in the literature. The method we developed addresses the limitations of these techniques by maintaining the 3D structure of the eye relative to the head and by being able to make clear distinctions between different relevant measurements.

As an alternative to microCT imaging, digital image registration techniques can also be used to reconstruct a volume from serial sections through an insect eye (Hung and Ibbotson 2014). Although a volume reconstructed from sections allows measurements to be made with histological resolution, section distortions and alignment errors may limit the accuracy of calculations on the 3D structure. The radius of curvature of a corneal transect can also be measured from a section (Schwarz et al. 2011) or an micrograph of a whole eye (Bergman and Rutowski 2016) and combined with a measurement of facet diameter to calculate the IO angle. When used on butterflies, this technique closely matches the results of the pseudopupil method (Bergman and Rutowski 2016). However, measuring the local radius from the eye’s profile only provides information along a 2D transect without describing the eye’s entire visual field, and calculating the IO angle in this way has similar limitations to our method in the presence of skewed CC (discussed further in the following section and the Supplemental Methods).

The pseudopupil technique allows IO angle (and facet diameter) to be measured directly from live insects and maintains the topology of these variables with reference to the visual world (Stavenga 1979). However, it is rarely applied across the entire FOV – the equipment’s geometry typically limits the measurement range to ±70° in azimuth and elevation (Rutowski et al. 2009) – and cannot measure the dimensions of internal eye structures that influence sensitivity; it is also difficult to use directly on compound eyes that have uniformly black iris pigment and lack a pseudopupil (as is the case for many hymenopteran eyes, including those of honeybees and bumblebees). Antidromic illumination is an alternative in such cases, whereby a light source placed inside an insect’s head to create a pseudopupil with light emerging from the eye’s facets along the receptors’ reverse optical path. This illumination method has been used to measure IO angles into the frontal visual field of several species (Kirschfeld 1973, Spaethe and Chittka 2003, Dyer et al. 2016), however, desert ants and honeybees are the only species where this method has been used to determine their full FOV and the IO angle topology (Seidl 1982, Zollikofer et al. 1995). The method we have developed to calculate the corneal FOV and IO angle can broadly be applied to any apposition compound eye regardless of its iris pigmentation and could even be used on preserved insects with intact corneas. This approach could also be used to determine the visual properties of superposition eyes (Land and Nilsson 2012), although modifications would be required to account for the larger optical effect of crystalline cones in this eye type. Unique to our approach is the ability to describe vision in world-referenced coordinates, which allows our findings to be quantitively compared to the results of studies on the vision of other species.

### Comparison to other studies on honeybee and bumblebee eyes

Honeybees have been the subject of many previous studies on eye anatomy and visual capabilities using many different techniques. As such, they are an important reference species for assessing the validity of the analysis method presented here. In addition, honeybees have no distinct size polymorphism (Streinzer et al. 2013), which improves the robustness of comparisons between different studies by minimising differences caused by eye size variation. As a case in point, the honeybees in our study and those analysed in Streinzer et al. (2013) were almost identical in body size (our specimens had ITWs of 2.9 mm and 3.0 mm vs. 2.9±0 mm from Streinzer et al. (2013)), and had similar eye surface areas (both 2.4 mm^2^ vs. 2.5±0.1 mm^2^), facet numbers (5440 and 5484 vs. 5375±143) and maximum facet sizes (25.4 *μ*m and 25.6 *μ*m vs. 25.2±0.3 *μ*m). Thus, our method is capable of describing corneal eye properties to within 5% of the values provided by replica-based techniques^4^. The thicknesses of eye components have previously been described from sections through a honeybee’s compound eye (Greiner et al. (2004); in the forwards facing areas of our honeybee eyes, we measured similar lens (34 *μ*m and 37 *μ*m vs. 28 *μ*m) and CC thicknesses (46 *μ*m and 50 *μ*m vs. 55 *μ*m). However, our retinal thickness measurements were substantially lower (219 *μ*m and 223 *μ*m vs. 320 *μ*m), and the value from histology lies above the range we measured for that parameter. Although our thickness measures are consistent between individuals, the relative difference between our measurement of retinal thickness and that of Greiner et al. (2004) is substantially larger than the error we estimated for measuring corneal eye properties. One likely explanation for this difference is that our definition of thickness is based on the local surface NV from the lens, and the distance until its intersects the upper surfaces of the CC, retina, and lamina interfaces (Fig. 2C), whereas Greiner et al. (2004) directly measured the rhabdom length from serial sections. Rhabdom length, rather than retinal thickness, is the relevant dimension for calculating optical sensitivity and, as a rhabdom may not necessarily lie perpendicular to the cornea, its length can evidentially exceed structural thickness measurements (the ratio by which it does so could also vary across the eye).

Previous studies on *Apis* have not calculated corneal IO angles, but several have used an antidromically illuminated pseudopupil to measure IO angles directly. Values for the average IO angle in a honeybee’s frontal visual field have previously been measured as 1.8° (Kirschfeld 1973) and 2.0° (Seidl 1982), with the latter recording 1.2° in the acute region. In the frontal and acute regions, the corneal IO angles calculated for our honeybees (frontal: 1.4° and 1.7°; acute: 0.9° and 1.0°) are lower than direct measurements from the IO angle but they exceed IO angle measured by Seidl (1982) in the dorsal and lateral regions (Table S2). The primary source of error in these measurements is that the honeybee’s ommatidial viewing axes are skewed from the corneal normal, and this misalignment varies across the eye (Stavenga 1979). This skewness is visible in sections through compound eyes (Baumgärtner 1928), and is unfortunately not correctable without segmenting the individual CC to use in optical modelling. As a result of this, our method underestimates the lower range of honeybee IO angles by up to 30% but could overestimate other angles by 30% to 60% (note that these errors will vary if the pattern of CC skewness differs in the compound eyes of other species). Because we calculate FOV directly from the corneal projection, we also underestimate the honeybees’ complete FOV as this can be enlarged through skewing the CC. An individual eye’s FOV is reported by Seidl and Kaiser (1981) as *nearly* hemispherical, while the corneal FOV of the eyes measured in this study span a quarter (25% and 27%) of the world sphere. The greatest differences between the FOV extents calculated by these methods appear to occur in the ventral and posterior regions; Seidl and Kaiser (1981) reported the honeybee FOV extends down to −90° in elevation and back to −156° in azimuth (at 0° el.), while we find the corneal FOV extends to −60° in elevation (at −60° az.) and to −107° in azimuth (at 0° el.). Additionally, Seidl and Kaiser (1981) found binocular overlap in the dorsoanterior, frontal, and ventroanterior regions, while our corneal FOV only indicated binocular overlap dorsoanteriorally. Given that pseudopupil measurements found a larger FOV and often larger IO angles than our corneal projection method, the optical axes of ommatidia must generally diverge in honeybee eyes to create a larger FOV at the expense of resolution. Although this demonstrates a limitation of our method, it highlights the importance of understanding CC orientation if detailed analyses and comparisons of visual fields are to be made within and across species.

The visual systems of bumblebees have received much less attention than those of honeybees. Nonetheless, there are several studies providing data against which we can compare our results. The body size of the two medium sized bumblebees in our study were similar to *B. terrestrìs* workers analysed by Streinzer and Spaethe (2014) (we measured an ITW of 4.0 mm for both bees, vs. 3.9±0.6 mm from Streinzer and Spaethe (2014)), as were their maximum facet sizes (25.0 *μ*m and 25.2 *μ*m vs. 25.1±1.9 *μ*m). However, the surface areas of our bee’s eyes were slightly smaller (2.2 mm^2^ and 2.4 mm^2^ vs. 2.8±0.6 mm^2^) and had fewer facets (4941 and 5505 vs. 5656±475). As both eye and body size vary substantially between bumblebee individuals, we did not estimate their ‘worst case’ errors as we did for honeybees. Nonetheless, the similarity in the majority of measurements between our study and that of Streinzer and Spaethe (2014) suggests good agreement between the different methods used to obtain them and provides further support for the validity of our method. The pseudopupil technique has also been used with antidromic illumination to measure the IO angles from the mediofrontal area of bumblebee eyes as function their body size (Spaethe and Chittka 2003), from small (ITW: 2.8 mm to 3.0 mm, mean IO angle: 1.5° from 6 bees) to medium sized individuals (4.0 mm to 4.2 mm, 1.2° from 4 bees). In comparison, we found relatively larger corneal IO angles (facing frontally) for our equivalently sized small (3.0 mm, 2.7°) and medium (both 4.0 mm, 2.2° and 2.4°) sized bees. However, our method shows that the corneal IO angle of bumblebees decreases from their frontal to lateral FOV (Fig. 5A), and by −45° az. the corneal IO angles (small bumblebee 1.5°, medium bumblebee 1.3°) closely match the measurements Spaethe and Chittka (2003) made using the pseudopupil method. The eye-referenced location of IO angle measurements by Spaethe and Chittka (2003) does not provide a clear world-referenced viewing direction and it is conceivable they were taken from eye regions directed somewhat laterally, in which case our results provide similar values. This highlights the importance of using a world-reference for visual studies, even when comparing between individuals of the same species. Regardless of eye size, we found the minimum corneal IO angles of both bee species were oriented somewhat laterally (Fig. 6Aii), challenging the common assumption that the frontal visual field is always most relevant for acute insect vision.

### *Allometry of* B. Terrestris *eye structure*

We found a scaling exponent of 0.45 for eye surface area vs. ITW for the *B. terrestris* workers in this study (Fig. 3A). This is substantially lower than for the between species allometry rate found across a range of 11 *Bombus* species (0.73) that had a similar range of body sizes (Streinzer and Spaethe 2014) indicating that the absolute investment in compound eyes varies more between *Bombus* species than between individual *B. terrestris*. Similar scaling exponents in other *Bombus* species would provide a clear indication that, independently of any factors influencing body size, visual requirements have influenced the evolution of eye size in different bumblebee species. We also calculated the scaling exponents of facet number and diameter as a function of eye size^5^ from the 11 *Bombus* species described by Streinzer and Spaethe (2014). We find that the number of facets has a higher scaling rate between *Bombus* species (1.39) than within *B. terrestris* (0.61). Conversely, the maximum size of facets had a lower scaling rate between species (0.26) than within *B. terrestris* (0.70). Without considering their visual topology, this indicates that the scaling rates of eye properties differ both between and within bumblebee species suggesting that the facet number, and thus resolution, is a species-specific adaptation. Although the maximum facet diameter is similar between species, substantial variation in facet diameter occurs between *B. terrestris* individuals of different sizes (Fig. 3E) (Spaethe et al. 2001). A recent study on the allometry of wood ant eyes from different colonies also showed that, despite maintaining similar scaling exponents for total eye size between colonies, two colonies invested in more facets as eye size increased, while another invested in larger facets (Perl and Niven 2016). Evidently, varying the parameters of eye allometry allows for substantial fine-tuning of the visual performance of individuals, both between and within species. The allometry of visual performance of bumblebees may influence which individuals and species can most effectively forage at specific floral resources (Dafni et al. 1997).

### *Allometry of B*. Terrestris *visual capabilities*

Within a species, variation between an individual’s visual capabilities could impact on their foraging ability. Behavioural studies investigating the influence of size on visual performance have shown that larger bumblebees do indeed i) have more acute vision when trained to discriminate visual targets (Spaethe and Chittka 2003), ii) are able to fly at lower light intensities (Kapustjanskij et al. 2007), and iii) are more efficient foragers (Spaethe and Weidenmüller 2002). Spaethe and Chittka (2003) also found that an increase in body size of 34% halved the minimum angular object size that a bee could identify, yet they noted that the allometric improvement in IO angle they measured could not directly predict the improvement in behavioural visual acuity. We applied the linear regression equation calculated by sSpaethe and Chittka (2003) to predict the minimum detectable object size for the small and medium bees in our study (ITWs 3.0 mm and 4.0 mm), giving visual angles of 8.4° and 5.2° respectively^6^ (a 38% decrease). While these angles are substantially larger than the corneal IO angles calculated using our method, the greatest relative improvement in IO angle (at any matching direction in the common FOV) between our small and medium sized bees is 34%, surprisingly close to the relative 38% improvement predicted by the behavioural study. This local acuity improvement is directed frontally (−9° el., −15° az.), a region of the visual field that is likely to be involved in target discrimination and that has been shown to be important for measuring optic flow for flight control (Baird et al. 2010, Linander et al. 2015). Our findings suggest that relative differences in acuity can indeed be accurately estimated from the allometry of the local IO angle in the relevant location of the visual field.

Interestingly, bigger eyes do not always provide increased visual performance – bumblebees with larger eyes do not exhibit differences when discriminating between different periodic patterns (Chakravarthi et al. 2016). Given that the IO angle on the mediofrontal eye area of medium sized bumblebees is 1.2° (Spaethe and Chittka 2003), relatively poor resolution limits have been measured for target detection (2.3°) (Dyer et al. 2008, Wertlen et al. 2008) and pattern discrimination (4.8 °/cycle) (Chakravarthi et al. 2016). The limits obtained from these behavioural experiments are approximately twice as high as what would be expected based on the sampling frequency (Snyder 1979). How the optics of the compound eye lenses focus light may also limit visual acuity as oversampling, where the optical cut-off frequency is lower than the sampling frequency, is an additional limiting factor that we have not considered here (Snyder 1979). Our analysis shows that lens thickness does vary across all bee’s eyes (Fig. S2B), which would result in local differences in focal length and may lead to topological variation in acceptance angle and, thus, the optical cut-off frequency. An approximately 25% increase in acceptance angle has indeed been found between frontally and laterally facing ommatidia in honeybees (Rigosi et al. 2017). Two additional confounds for determining visual acuity from anatomical measurements are that acceptance angle varies between states of light and dark adaptation (Warrant and McIntyre 1993), and that both object illumination and contrast also influence an eye’s effective resolution (Snyder et al. 1977, Warrant 1999).

### *Allometry of B*. Terrestris *visual fields*

In this study, we found that the corneal visual field of bumblebee eyes increased with eye size (Fig. 3C). This was not simply a consequence of having a larger eye, but also of a change in eye shape such that the surface was projected onto a larger angular area. We also identified an area of binocular overlap not previously reported in bumblebees. The extent of this corneal binocular overlap, directed both frontally and dorsofrontally (Fig. 4C), increased rapidly with body size. Bumblebee workers have been found to approach artificial (Reber et al. 2016) and natural flowers (Orth and Waddington 1997) from below, which would place the visual target dorsofrontally, a region where we also found that corneal IO angle decreases with eye size (Fig. 7A). Hence, larger bumblebees would view the flowers they approach with a more acute and larger binocular visual field, which would potentially improve their visual discrimination or control of landing relative to smaller bees. Such potential benefits of the binocular overlap would be an interesting topic of further behavioural investigations.

Surprisingly, the scaling exponent (as a function of body size) found here for bumblebees’ corneal FOV is nearly identical to that found from the optically measured FOV of differently-sized butterfly species (Rutowski et al. 2009). This is the case both for the FOV of a single eye (we found 0.17 vs. 0.14) and the binocular FOV (0.79 vs. 0.82). In contrast, the FOV of desert ants remains similar despite a nearly two-fold increase in head size (Zollikofer et al. 1995). Given the common scaling rates between the FOV of *B. terrestris* workers and butterflies, we hypothesise that increasing the visual field of each eye, and the binocular overlap between eyes, at the identified rates may be a general strategy of compound eye enlargement among different groups of *flying* insects. While visual field extent has rarely been considered by previous studies on insect vision, increasing FOV size has been shown to improve the performance of visually-guided behaviours such as navigation (Wystrach et al. 2016) and visual egomotion detection (Borst and Egelhaaf 1989), and is likely to improve the performance of larger bumblebees performing these visually guided behaviours.

### *Allometry of B*. Terrestris *visual topology*

Local variation in the scaling rate of eye properties will cause eye-dependent variation in the topology of visual capabilities. The region with the lowest corneal IO angle scaling exponent (leading to improved visual resolution) is directed dorsofrontally (Fig. 7A), while a positive but relatively similar scaling exponent for facet size occurs across the visual field (Fig. 7B). In contrast, the scaling rate of IO angles vs. body size of Orange Sulphur butterflies was strongest in the ventral, anterioventral, and dorsal eye areas, while their facet diameters were also found to increase at a uniform rate across the areas measured (Merry et al. 2006). Again, the results from ants are qualitatively different from the results for bees and butterflies: the scaling exponent of facet diameter varied between the eye areas of wood ants, being highest in the dorsal and anterior areas (Perl and Niven 2016), while a study on desert ants found IO angle scaled similarly between lateral and dorsal eye areas (Zollikofer et al. 1995). Pseudopupil measurements along a vertical transect of the eyes of damselfly species found that the maximum diameters and minimum IO angles were both influenced by eye size and habitat (Scales and Butler 2016). Although the scaling exponents along the eye transects were not measured, damselflies living in dim, cluttered habitats also appeared to have more prominent eye specializations than those living in open habitats, independent of eye size.

To our knowledge, this is the *first* study to investigate the 3D topology of retinal thickness in insects (Fig. 5) and it is evident from our analysis that retinal thickness varies substantially across all bee eyes. If differences in retinal thickness are translated into equivalent differences in rhabdom length, this would influence optical sensitivity across each eye by 20% to 50% for bumblebees and 53% for honeybees^7^. Retinal thickness is typically highest ventrally and posteriorly (Fig. 5B), where higher retinal sensitivity may compensate for the reduced effective aperture resulting by the skewed CC in these regions (Stavenga 1979). Retinal thickness has a positive scaling exponent across the majority of the visual field (Fig. 7C), which would improve the optical sensitivity of larger bees. Unexpectedly, we identified that retinal thickness increases at a higher rate in the dorsal hemisphere (Fig. 7C) and would provide larger bumblebees with relatively increased dorsal sensitivity that may be useful to visually identify downwards facing flowers that are not directly illuminated by sunlight (Makino and Thomson 2012, Foster et al. 2014). Our results demonstrate that, in addition to facet size, retinal dimensions offer substantial scope for insects to fine-tune optical sensitivity across their visual fields, a point that appears to have been overlooked by previous studies.

The visual topologies and scaling exponents we measured for bumblebees partially support our initial hypothesis that the increased resources of a larger eye would primarily be invested in improving the capabilities of a small visual region. The improved visual resolution of larger bees is primarily directed dorsofrontally, but the scaling of facet diameter and retinal thickness would lead to increased optical sensitivity across their entire field of view. As a result, we now hypothesise that, for a given insect group, specific regions in the visual field may have certain ‘ideal’ requirements for resolution and/or sensitivity based on the visual information available in their specific habitat and their behavioural ecology. Once the size of an eye allows such a threshold to be reached, additional area could then be invested in improving the visual capabilities of other regions. This revised hypothesis incorporates our findings that differential allometry of *Bombus* eye properties flexibly allow their visual capabilities to be improved either locally or globally across their changing FOV.

### Conclusion

Analysing the 3D structure of insect eyes to determine a holistic description of their visual capabilities provides insight into how the morphology of eyes have evolved to sample visual information from the world. We find that differential scaling of the morphology between eye areas allows *bigger bumblebees* to invest the increased resources of a larger eye in *improved sensitivity* across an enlarged visual field. Yet, studying the allometry of their entire visual topology also indicated specific regions with a high investment rate that may have otherwise been overlooked, such as the dorsofrontal region of both *enlarging binocularity and increasing resolution*, or the high rate of thickening in the dorsal facing retina. Important visual information is presumably viewed by bumblebees in these regions of their visual fields, indicating a promising avenue for further behavioural experiments, such as by using virtual-reality to manipulate the visual cues at specific regions in an insect’s FOV (Stowers et al. 2017) and observational studies to identify what bumblebees view in those regions when flying through natural environments (Stürzl et al. 2015). The differential allometry between eye areas undoubtedly enables insects with larger eyes to improve the quality and reliability of visual information by having a better capacity to match their visual capabilities to the requirements of both the environment and behaviour.

## Acknowledgment

We would like to thank Carina Rasmussen, Eva Landgren, and Ola Gustafsson for facilitating sample preparation and providing access to the Microscopy Facility at the Department of Biology, Lund University. We are also grateful to Karin Odlén, Julia Källberg, Per Alftrén, and particularly Viktor Håkansson, for assistance preparing samples and analysing data, and also to Viktor for diligently managing tomographic reconstructions during our beamtime. Our imaging was performed at Diamond Light Source (proposals 13848 and 16052), where we were pleased to receive assistance from Qiang Tao, David Wilby, Rajmund Mokso, and Kazimir Wanelik. Gavin Taylor is thankful to have received a stipend from Carl Tryggers Stiftelse (CTS15:38) and an endowment from the Royal Physiographic Society of Lund, while Emily Baird acknowledges financial support from the Air Force Office of Scientific Research (FA8655-12-1-2136), the Swedish Research Council (2014-4762), and the Lund University Natural Sciences Faculty. Pierre Tichit received funding by Interreg Project LU-011, and Marie Schmid received funding from the Erasmus+ program.

## Supplemental material

**List of supplemental abbreviations**
az.: Azimuth
BP: Border point
CC: Crystalline cone
CLP: Corneal linkage point (of a facet dimension measurement)
D: Facet diameter
el.: Elevation
EV: Eye volume
FOV: Field of view
IDW: Inverse distance weighted
ITW: Inter-tegula width
IO: Inter-ommatidial
NV: Normal vector
R: Radius of curvature
SP: Sampling point
WP: World point
*ΔΦ*: IO angle
*σ*: Ommatidial axis density

## Supplemental figures

**Figure. S1:**
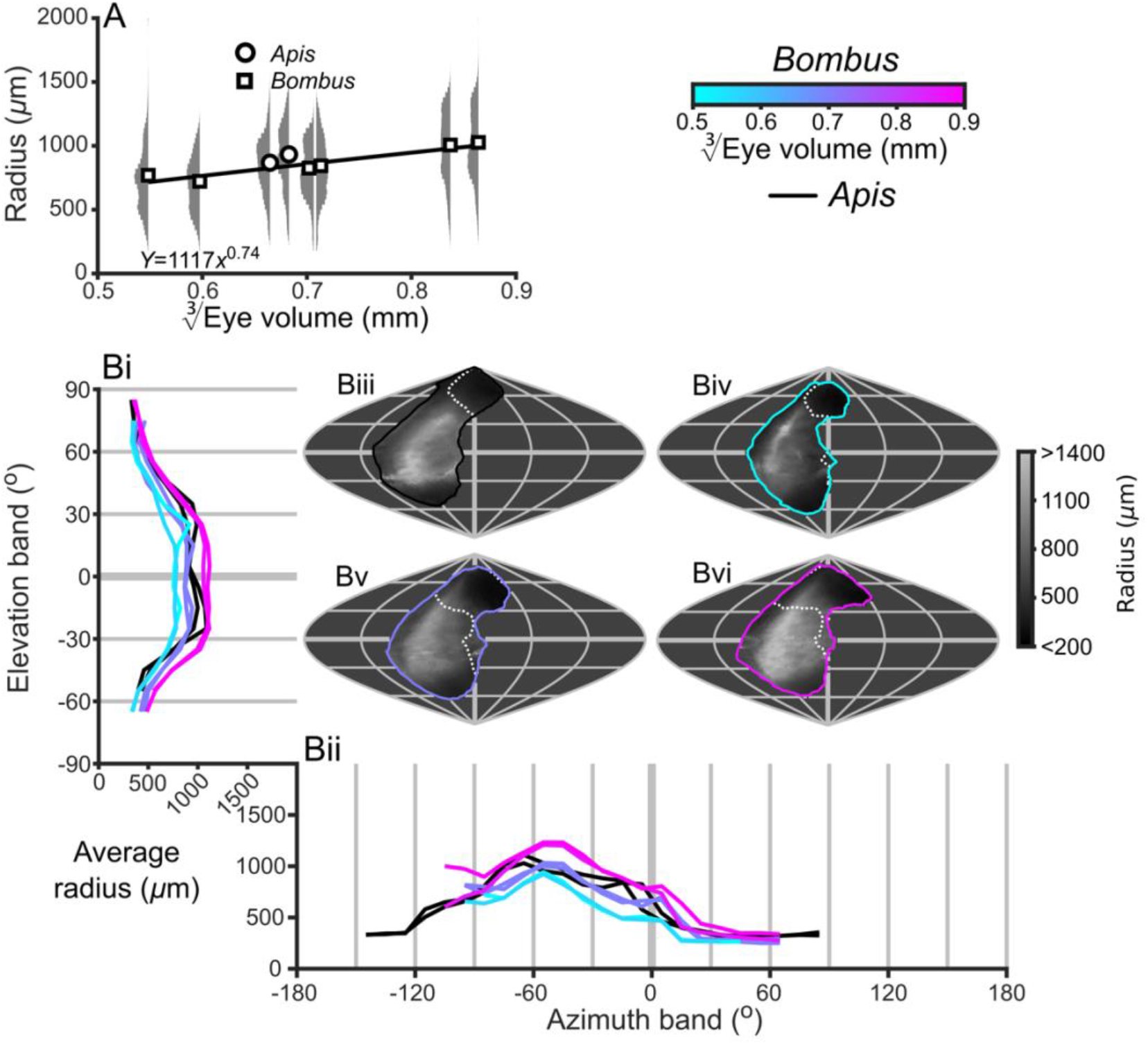
Description of the local radius of curvature of compound eyes. **A**, the average radii and relative distribution for each bee vs. 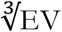. The plotted curve results from fitting the function parameters to the measurements for *Bombus* (additional details in Table S1). **B**, the topographic distribution of *R* (indicated by the grey colour bar) projected from each bee into the visual world, shown as sinusoidal projections for a honeybee (iii), and small (iv), medium (v), and large (vi) sized bumblebees (azimuth and elevation lines are plotted at 60° and 30° intervals respectively). The average *R* profiles are shown for elevation (i, *R* is averaged across all azimuth points in each eye’s FOV as function of elevation) and azimuth (ii, *R* is averaged across all elevation points in each eye’s FOV as a function of azimuth). The white dotted line across the topology in Biii to Bvi indicate the border of the bee’s corneal binocular FOV. The cyan-to-magenta colour bar indicates 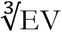 of *Bombus* for each curve in all panels, while black lines indicates honeybees.

**Figure. S2:**
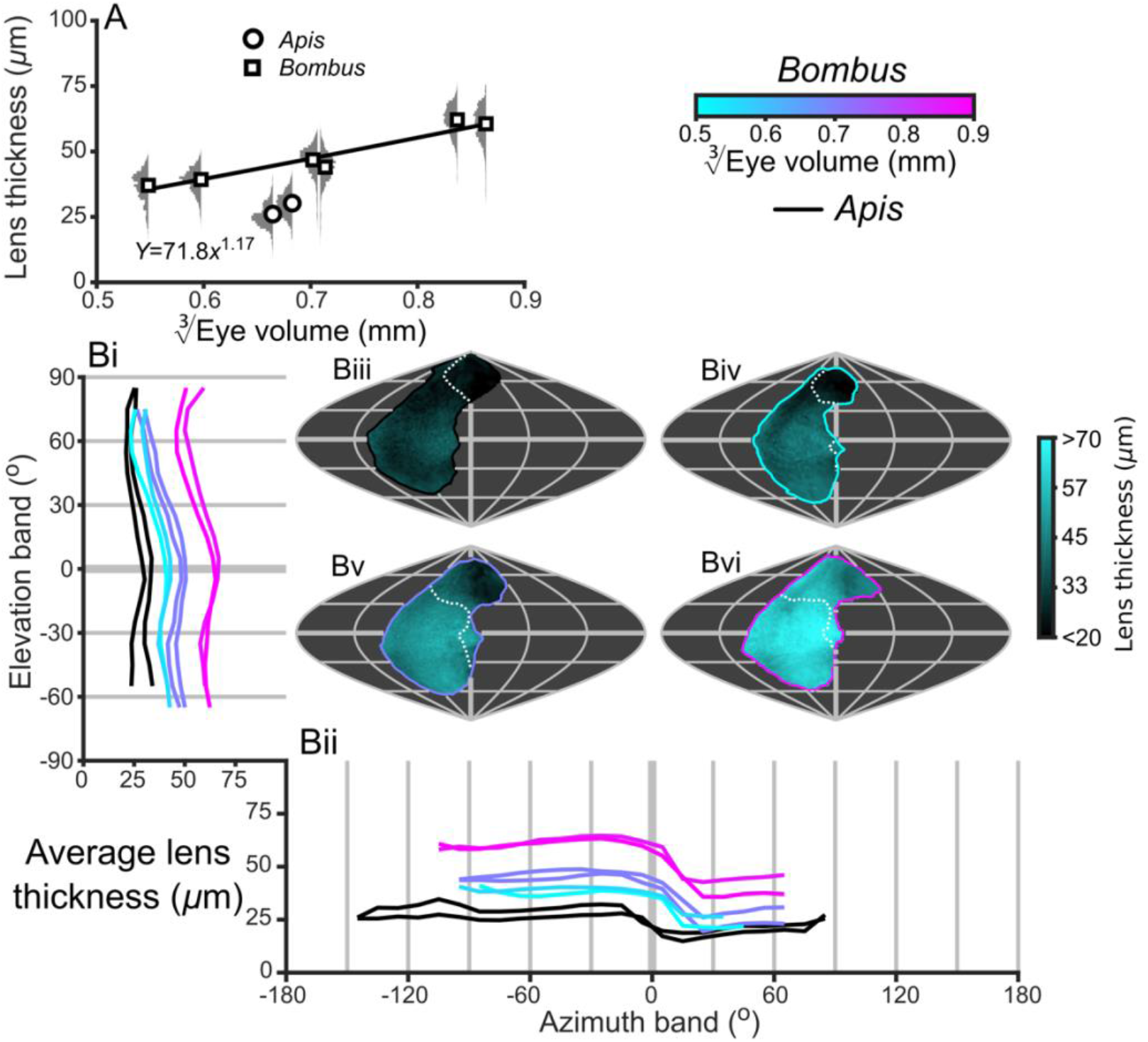
Description of the local lens thickness of compound eyes. **A**, the average lens thicknesses and relative distribution for each bee vs. 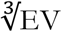. The plotted curve results from fitting the function parameters to the measurements for *Bombus* (additional details in Table S1). **B**, the topographic distribution of lens thickness (indicated by the cyan colour bar) projected from each bee into the visual world, shown as sinusoidal projections for a honeybee (iii), and small (iv), medium (v), and large (vi) sized bumblebees (azimuth and elevation lines are plotted at 60° and 30° intervals respectively). The average lens thickness profiles are shown for elevation (i, thickness is averaged across all azimuth points in each eye’s FOV as function of elevation) and azimuth (ii, thickness is averaged across all elevation points in each eye’s FOV as a function of azimuth). The white dotted line across the topology in Biii to Bvi indicate the border of the bee’s corneal binocular FOV. The cyan-to-magenta colour bar indicates 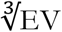 of *Bombus* for each curve in all panels, while black lines indicate honeybees.

**Figure. S3:**
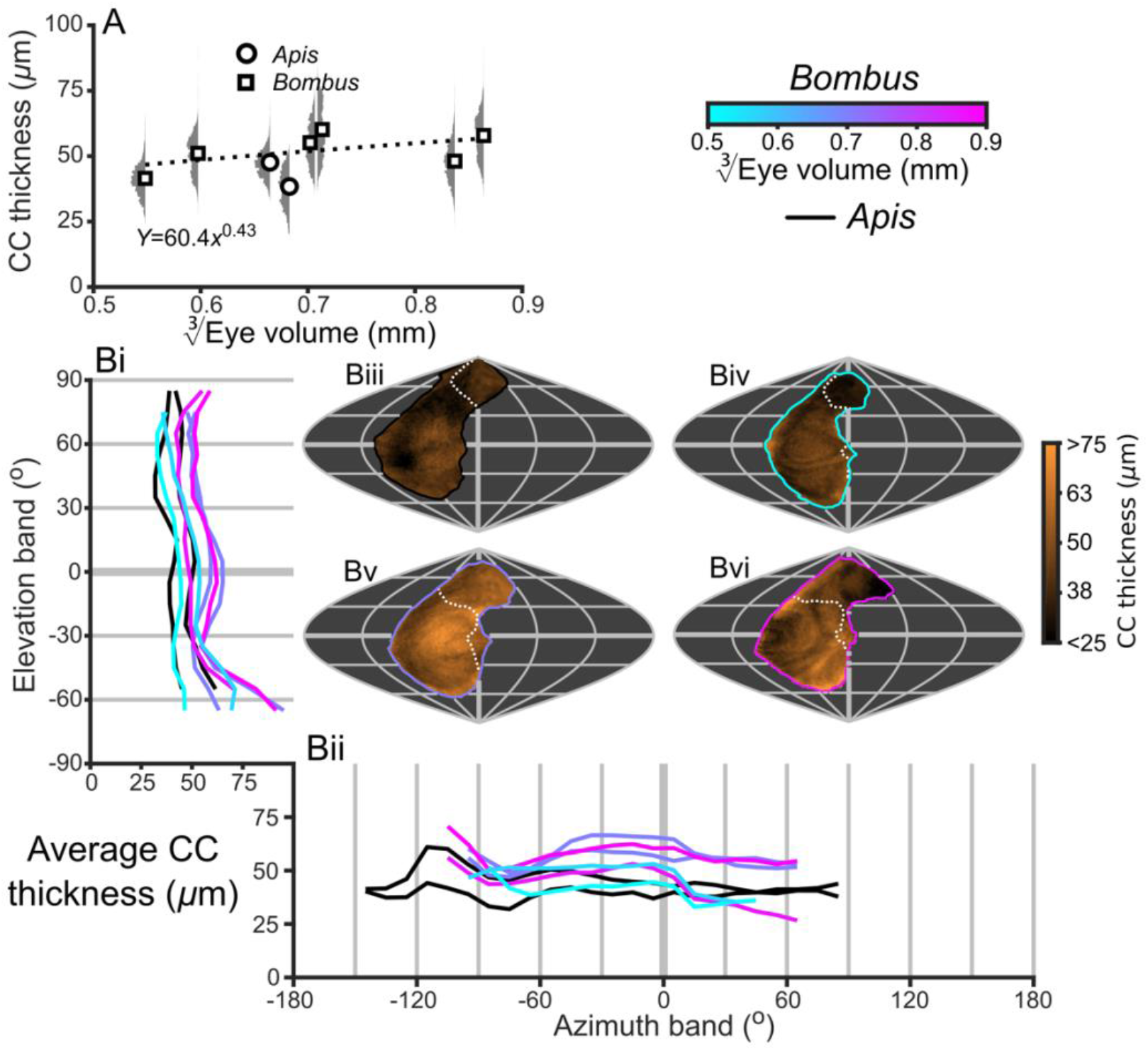
Description of the local CC thickness of compound eyes. **A**, the average CC thicknesses and relative distribution for each bee vs. 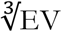. The plotted curve results from fitting the function parameters to the measurements for *Bombus* (additional details in Table S1), however, the correlation between CC thickness and 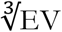 is not significant. **B**, the topographic distribution of CC thickness (indicated by the orange colour bar) projected from each bee into the visual world, shown as sinusoidal projections for a honeybee (iii), and small (iv), medium (v), and large (vi) sized bumblebees (azimuth and elevation lines are plotted at 60° and 30° intervals respectively). The average CC thickness profiles are shown for elevation (i, thickness is averaged across all azimuth points in each eye’s FOV as function of elevation) and azimuth (ii, thickness is averaged across all elevation points in each eye’s FOV as a function of azimuth). The white dotted line across the topology in Biii to Bvi indicate the border of the bee’s corneal binocular FOV. The cyan-to-magenta colour bar indicates 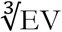 of *Bombus* for each curve in all panels, while black lines indicate honeybees.

**Figure. S4:**
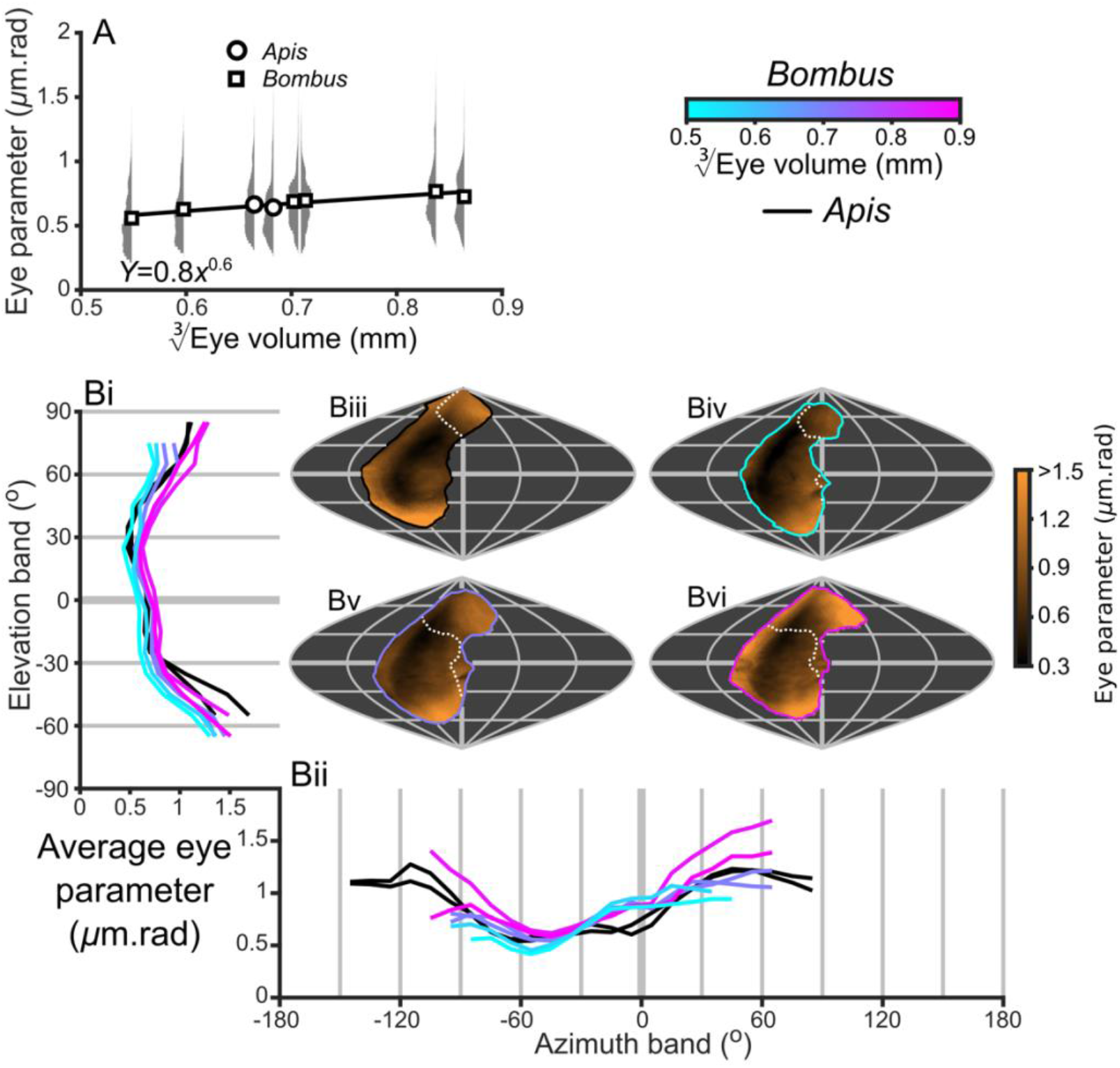
Description of the local eye parameter of compound eyes. **A**, the average eye parameters and relative distribution for each bee vs. 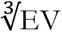. The plotted curve results from fitting the function parameters to the measurements for *Bombus* (additional details in Table S1). **B**, the topographic distribution of eye parameter (indicated by the orange colour bar) projected from each bee into the visual world, shown as sinusoidal projections for a honeybee (iii), and small (iv), medium (v), and large (vi) sized bumblebees (azimuth and elevation lines are plotted at 60° and 30° intervals respectively). The average eye parameter profiles are shown for elevation (i, eye parameter is averaged across all azimuth points in each eye’s FOV as function of elevation) and azimuth (ii, eye parameter is averaged across all elevation points in each eye’s FOV as a function of azimuth). The white dotted line across the topology in Biii to Bvi indicate the border of the bee’s corneal binocular FOV. The cyan-to-magenta colour bar indicates 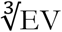 of *Bombus* for each curve in all panels, while black lines indicate honeybees.

**Figure S5:**
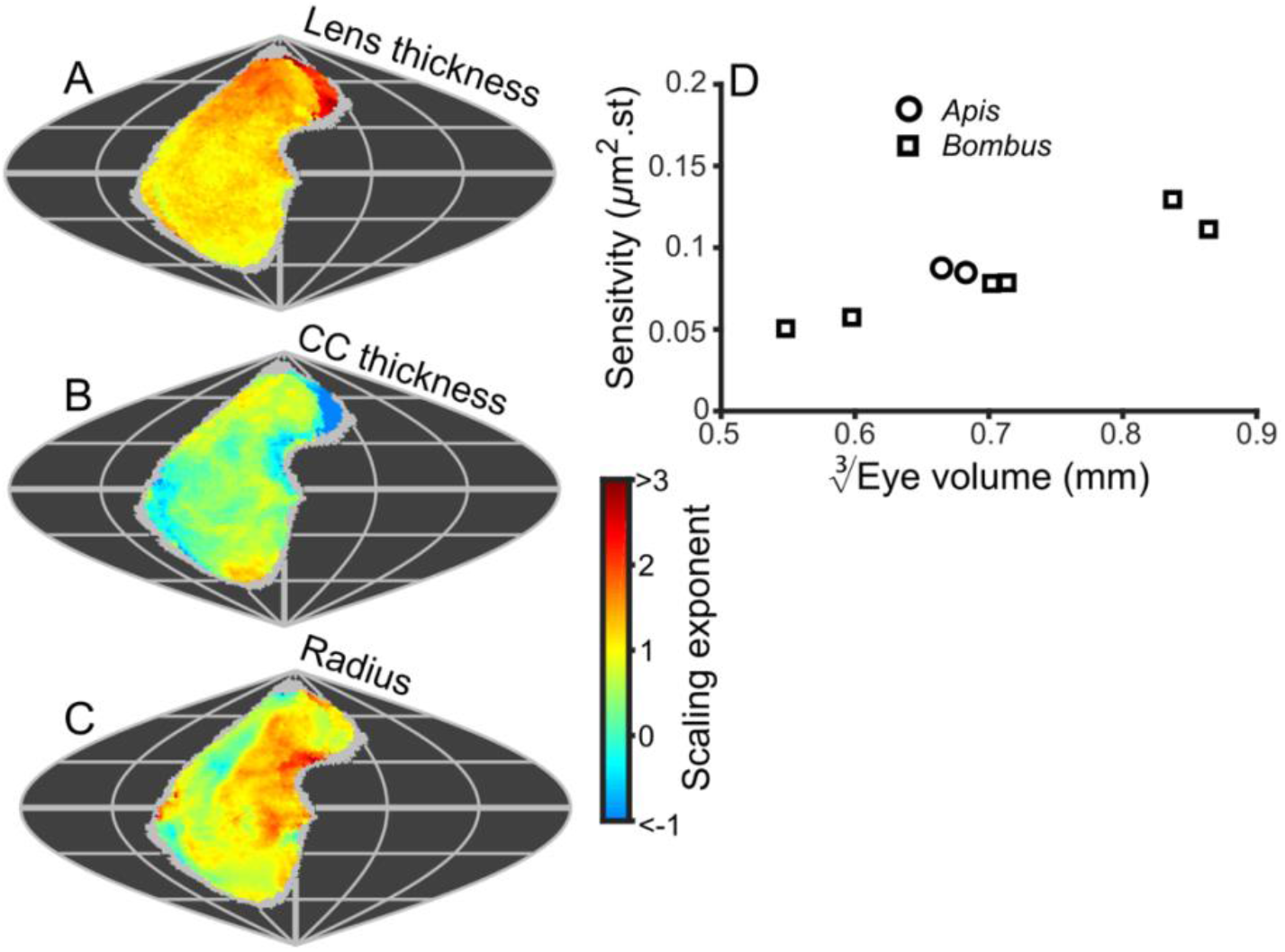
Scaling and sensitivity. **A** to **B** show maps of the scaling exponents calculated for bumblebees from the variable specified on each map. The scaling exponents for all variables use the blue-to-red colour bar. Positive scaling exponents (red) indicates the lens thickness (B), CC thickness (C), and radius of curvature (D), indicate a variable increasing with eye size. We limited the scaling exponent calculations to regions viewed by at least four bees, and the grey fringe in each map indicates the additional regions (viewed by three or fewer individuals) that were not used in the calculations. **D**, the average optical sensitivity calculated for each bee vs. 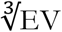. Note that this calculation assumes that the acceptance angle of each facet equals its corneal IO angle, and that the rhabdom length equals the retinal thickness.

**Figure S6:**
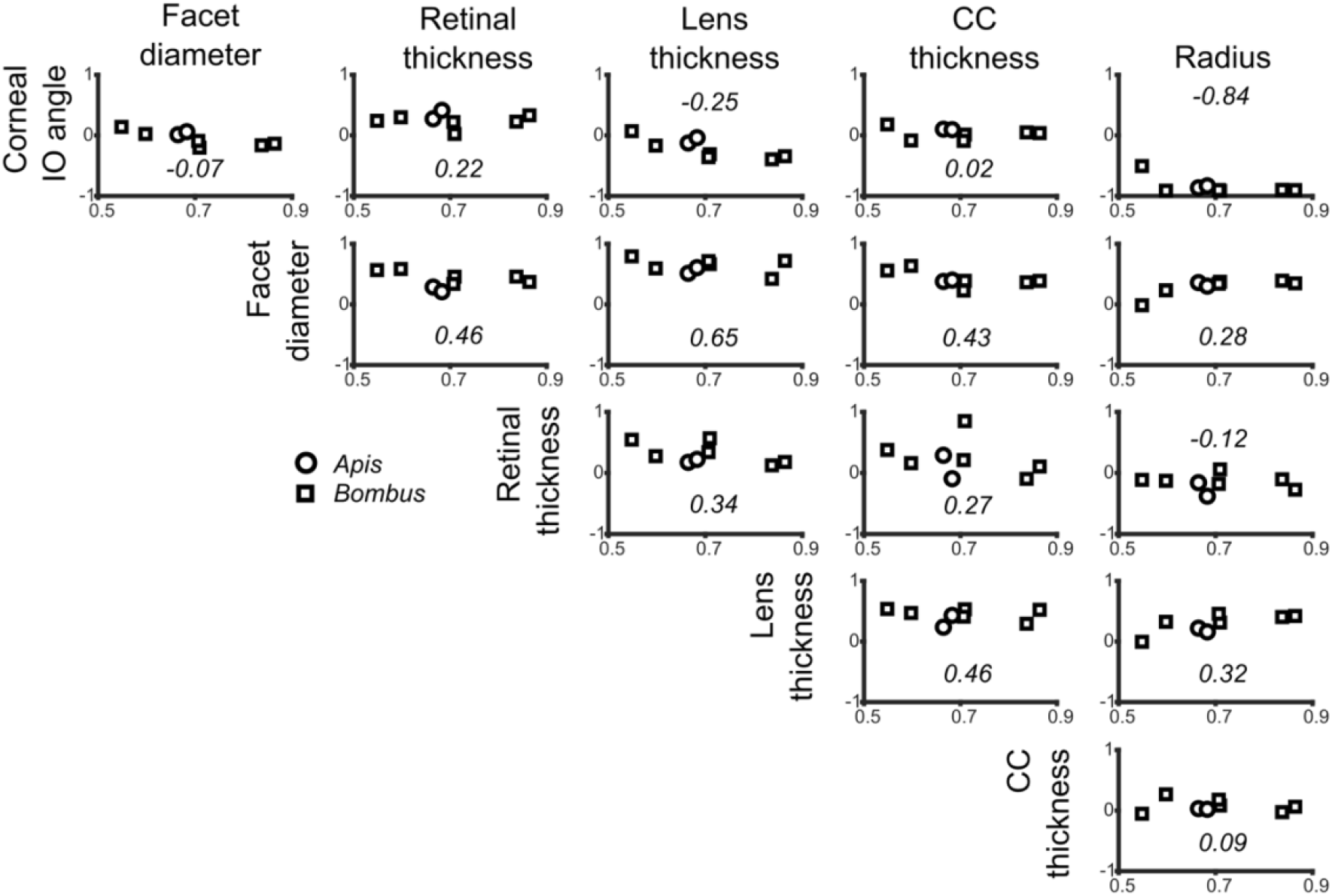
Correlation of local variable values. The matrix of graphs representing the correlation coefficient (*r*) between two local variables, measured for each bee vs. its 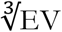. The vertically oriented variable name for each row are cross-referenced against the horizontally oriented variable name for each column to determine the correlation shown in each plot. The italicized number in each plot represents the average of that plot’s correlation coefficient for all bees.

## Supplemental tables

**Table S1:**
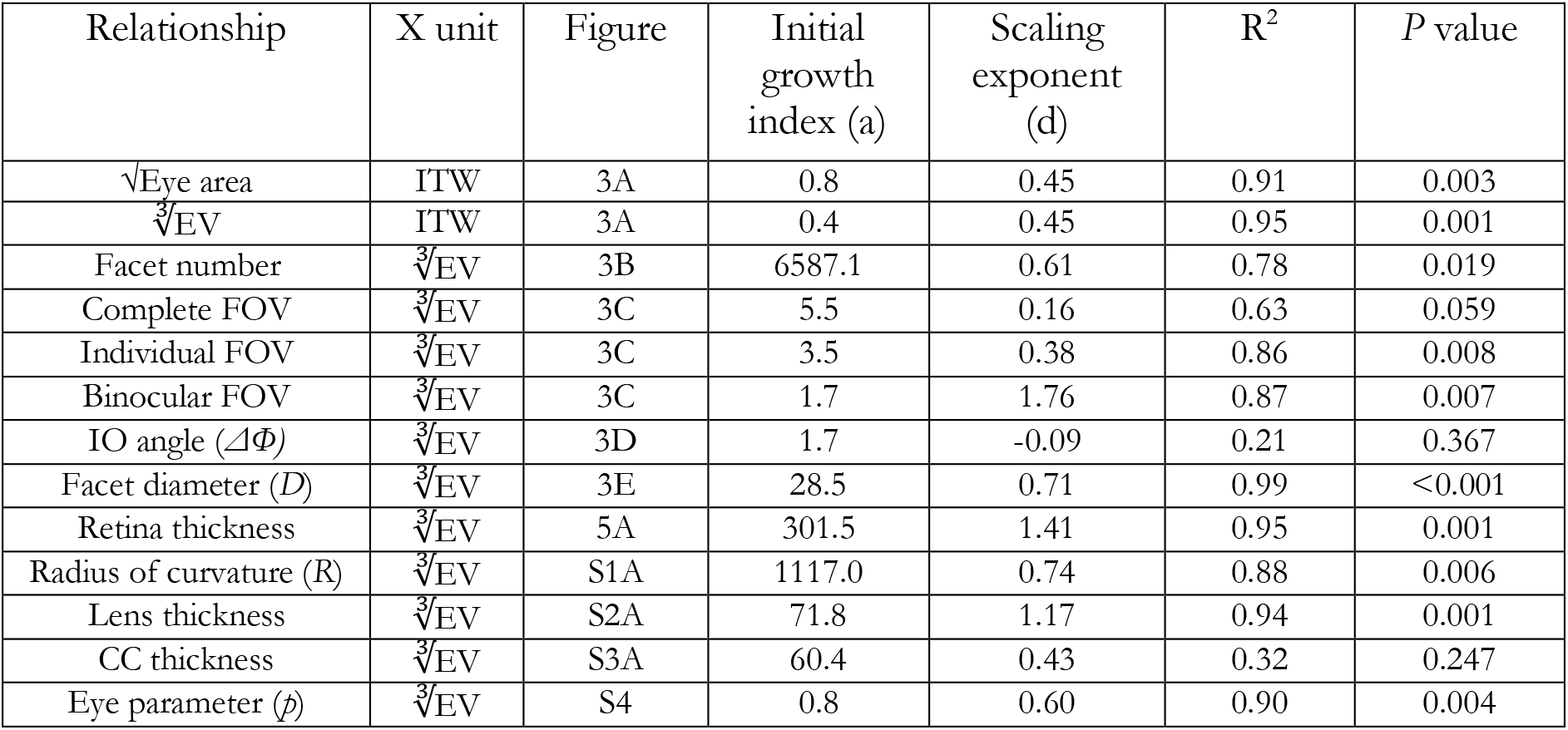
Parameters for the power functions calculated for bumblebees in this study. Note that non-significant correlations are plotted with dotted lines in the relevant figures.

**Table S2:**
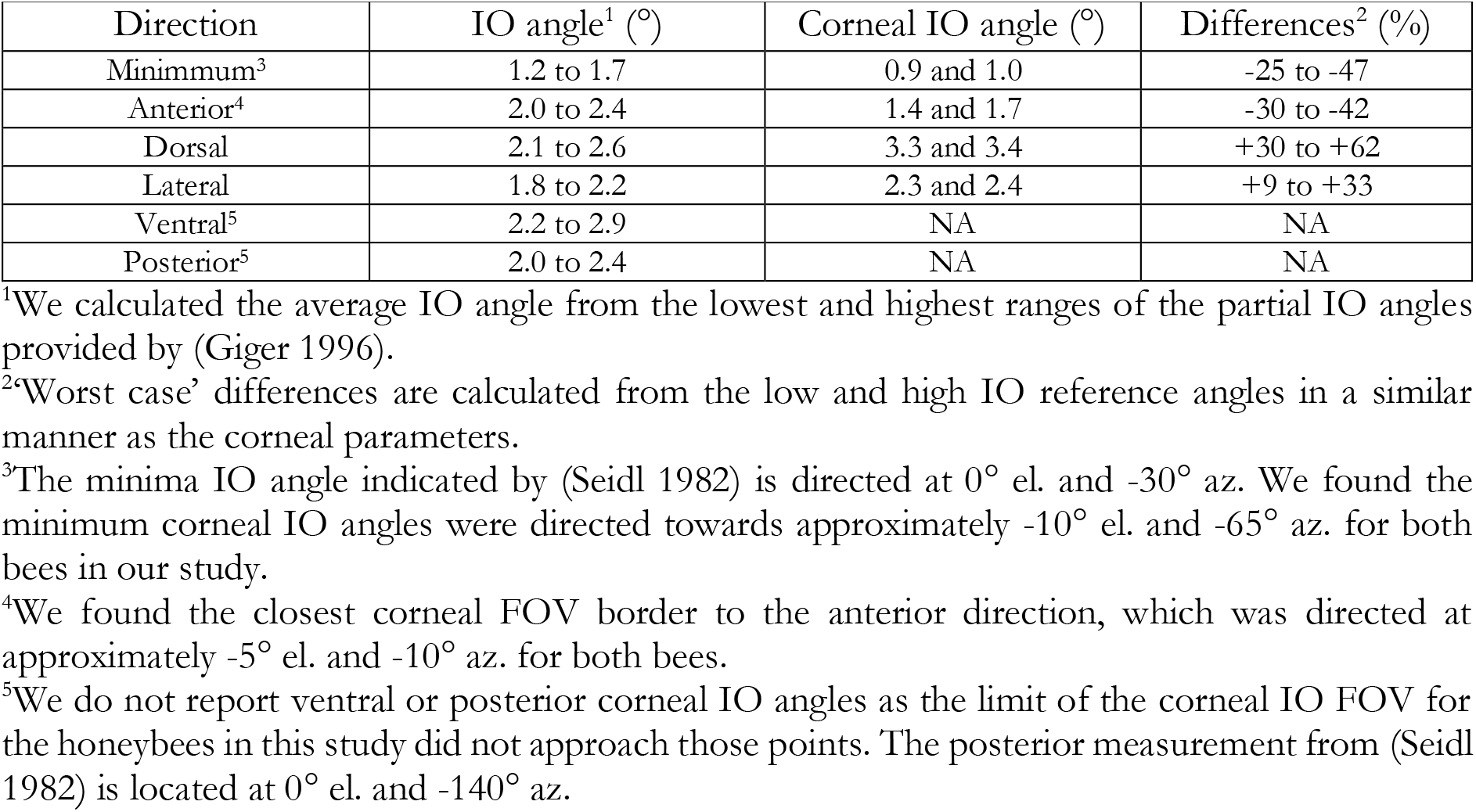
Topological differences between reported IO angle and our measured corneal IO angles for honeybees. We obtained the IO angles measured by Seidl (1982) from Fig. 2.2 in Giger (1996).

## Supplemental methods

### Volumetric analysis

Labelling was performed by using the ‘brush’ tool in the Amira Segmentation Editor to manually delineate the borders of the three gross anatomical structures in a compound eye (lenses, crystalline cones (CC), and retina; Fig. 1B,C) from individual slices through the greyscale volume. Features were labelled at 5 to 10 slices intervals, following which an interpolation tool was applied between the labelled slices. As high contrast existed between the exterior of the lenses and the surrounding air, this border could be automatically delineated using a threshold tool. Following manual segmentation and interpolation, the 3D label volumes were smoothed. The interface between the lenses-to-exterior, lenses-to-CC, and CC-to-retina were selectively labelled by using morphological operations between each pair of labels. The interface between the retina and lamina was identified by selectively rendering the volume immediately proximal to the labelled retina, and also viewing a surface generated from this label. The surface path selector was then used to trace a closed outline of this interface on the vertices of the retinal surface; the path could then be filled to select all enclosed vertices, which were converted to labelled voxels in the main label volume. The label volume was saved as a series of images for later analysis in Matlab.

Alignment of labelled compound eyes onto a head of the same species was performed manually in the Amira Project Viewer (Fig. 2A). A single head was used to align all compound eye replicates of a given species, and its greyscale volume was initially loaded into Amira and oriented such that the front of the head faced forwards and the tops of the eyes were parallel to the global coordinates of Amira. The greyscale volume of each labelled left compound eye was loaded, and either an isosurface or a volume rendering was used to visualize both the head and compound eye, depending on which visualization method provided the clearest representation of its structure. Each eye was then positioned and oriented such that it overlaid the matching compound eye on the head, which also required isometrically re-scaling the head for bumblebee eyes, as a single head from a medium sized bee was a used as a reference for the six compound eyes taken from differentially sized individuals. This re-scaling implicitly assumed that the relative position of compound eyes on the head capsule did not vary between bees with difference body sizes. Once aligned, the digital volume was resampled into the world coordinate frame, and then flipped about its x-axis (equivalent to mirroring about the sagittal plane of the head), the flipped eye was then positioned and aligned onto the head’s right eye. The affine transformation matrices for the head, and both the original and mirrored eyes were then recorded from the Amira console.

Facet dimensions were measured on compound eyes that were visualized using either an isosurface or volume rendering. Although the borders of individual facets were rarely visible in individual slices through the volume, these surface visualization techniques typically showed the fine borders between the facets. However, not all facets were visible on a given eye because the unavoidable presence of unpeeled glue or dust on the eye that prevented the visualization of some regions. To measure the size of a given facet, points were selected at the centre of the furthest sides of neighbouring ommatidia. Points were selected in pairs, such that each represented a line passing across the middle of the central facet to link the opposing sides of two neighbouring ommatidia (Fig. 2B). Twenty or more measurements were taken on any individual eye (depending on its size), and we aimed to distribute these evenly across the eye. We avoided including highly irregular facets (i.e. where one hexagon side length was much shorter than the others) in the analysis. The 3D coordinates of each selected point were saved in spreadsheets for importing into Matlab.

**Upon acceptance of this manuscript, we will upload the original data and volumetric label data for each eye to the Morphosource repository and provide Ascension numbers to access these data**.

### Calculation of global parameters

Data saved from Amira (labelled volumes, transforms, and facet measurements) were imported into Matlab with metadata for each bee (species, body size, etc.). After loading the labels, the eye volume (EV) was calculated from the total number of labelled voxels (including the lens, CC, retina and lamina contact labels, and their interfaces). We found that the most accurate method of measuring the total eye surface area was to take half the surface area of an isosurface fit around the corneal surface voxels.

### Transformation and alignment

The coordinates of the eye were transformed from volume coordinates into the head coordinate frame using the recorded transform, and mirrored to produce the right eye, which was also transformed appropriately. The three principle axes were then calculated from the corneal surface coordinates of both eyes, and both eyes were rotated such that the corneal surface were symmetric in the sagittal (roll) and transverse (yaw) planes. The vertical axes of the eyes (representing the coronal (pitch) plane) was then aligned to the world’s vertical axis.

### Facet shape determination

The six points measured for every facet were transformed into the same coordinate frame as the head, and the centre of the points measured for each facet was linked to the closest point on the corneal surface, termed the corneal linkage point (CLP). Locally, the corneal surface was treated as a flat (2D) plane on which the hexagonal facets were arranged. However, the measurements points of each facet were in 3D coordinates and were oriented in different directions, so the world’s azimuth and elevation directions were chosen as a 2D coordinate system to which all facets were aligned. To facilitate this, the normal vector (NV) was calculated at the CLP for each facet and the measurement points were rotated such that this vector pointed forwards. Each pair of measured points represented a line between the opposing sides of three adjacent facets, and we reduced the length of each line by 2/3 such that its ends represented the centre points of the outer two facets in a trio (assuming those facet centres were equidistant from the centre of the central facet). Thus, for each measured facet, we had 7 points representing its own centre and the centre of each neighbouring facet. We used this information to compute a Voronoi diagram, from which the central cell indicated the corner points of the measured facet (Fig. 2B). Although the corner points of each facet could have been measured directly, our method effectively averages the shape of the 7 facets covered by the initial six measurement points and is similar to the method of averaging facet diameter (D) over a row of five facets (Streinzer et al. 2013, Streinzer and Spaethe 2014, Streinzer et al. 2016). The area of the resulting hexagon was then calculated.

### Sampling variables on eye

Sampling points (SP) were selected across the 3D corneal surface of each eye at approximately equally spaced at 25 *μ*m intervals and formed the basis for eye-centric calculations. At each SP, we calculated the NV of the corneal surface and the coordinates of its intersection on a distant sphere that represented the visual world (Fig. 2C). The direction of each NV was then reversed to calculate the volume thicknesses at a given SP (Fig. 2C). We calculated the distance from the SP on the cornea to the vector’s intersection point with the lens-to-CC (L-CC) interface (local lens thickness), and then between the L-CC intersection and the subsequent intersection with the CC-to-retina (CC-R) interface (local CC thickness). If the vector subsequently intersected the retina-to-lamina (R-La) interface we calculated the distance to that intersection from the CC-R intersection (local retina thickness). If the vector did not intersect the R-LA interface, we instead found the closest voxel on the R-La interface to the CC-R intersection and calculated the distance between these two points as the retina thickness. We also calculated the NVs and sphere intersections from all voxels around rim of the corneal surface (where the cornea bordered the surrounding cuticle), although we did not calculate thicknesses for these border points (BP).

### Normal vector calculations from voxelized surfaces

We determined the corneal NV at each SP by selecting all corneal surface voxels within a local radius around the SP and fitting a 2^nd^ order polynomial surface to the selected voxels, and then calculated the NV from the derivative of the surface at the SP (Taylor et al. 2016). This method of determining a surface’s NV was implemented in a similar manner to the ‘coordinate transform’ method of parameterizing discretised surfaces used in computer vision applications (Stokely and Wu 1992). The choice of local radius increased or decreased the number of points included in the surface fitting, which influenced the NV and caused noticeable variation in the calculated corneal inter-ommatidial (IO) angle, as this was the result of finding the angle between multiple NVs (Fig. 2C). An iterative procedure is described in a following section which allowed us to choose an appropriate local radius for NV calculations on each eye. We also noticed that particularly flat corneal surfaces lead to inaccurate normal fitting, as information on the surface’s shape was lost when discretising the eye into voxels. To rectify this problem, we tested whether the outermost voxels in each selection varied in height with respect to the central voxels; if no variation was present (i.e. a flat plane of voxels had been selected), we enlarged the local radius until height variation was present across the selected points.

### Interpolation of hexagons between sampling points

The hexagon parameters at each SP were interpolated from the sparser set of CLPs using inverse distance weighted (IDW) interpolation (with *p*=3), where distances between points were based on the geodesic path lengths over the corneal surface (Shepard 1968). We devised a shape interpolation method based on the transformation vectors between the closest corner points of two centred hexagons. The corner points if the hexagon *CLP2* could then be described from the original hexagon’s corner points (CLP*1*) plus the transform (*T*), as CLP*2*=CLP*1*+*T*. As the facet shape determined for each CLP was initially rotated to face forwards, all corner points lay on the same plane (allowing 2D transformations to be used) and each facet’s alignment with respect to the world’s azimuth and elevation was preserved. These hexagon transformations were calculated between every pairwise combination of CLPs, and used to interpolate the shape of a hexagon at a given SP in the following manner:

- For a given SP only nearby CLP pairs were used in the interpolation (both were limited to be 100 *μ*m distant).
- For every pair of CLPs, the ratio of the distance between them and the SP was used to determine the magnitude of the transformation applied to CLP1. That is, the predicted hexagon (*H*) was, *H*=CLP*1*+*T*×*G*_SP→CLP*2*_/(*G*_SP→CLP*1*_+*G*_SP→CLP*2*_), where *G* denotes the geodesic path length between the two specified points. Hence, if the SP was closer to CLP*1, H* would be similar to CLP*1*, or vice versa if the SP was closer to CLP2.
- Predicting a hexagon from *N* nearby CLP pairs produced 6*N* corner points for a given SP. These were divided into six groups using the K-means clustering algorithm.
- The (weighted) average coordinate of each group then provided the six corner points of the interpolated hexagon at a given SP. The weight for each predicted hexagon was the inverse of the average distance for both CLPs to the SP.
- The area of the interpolated hexagon was calculated, as was the distance and angle to the six adjacent facet centres based on treating the interpolated hexagon as cell in a Voronoi diagram.

IDW interpolation was then used to directly predict the expected local facet surface area at each SP from the calculated facet areas of the CLP. On average, the expected facet surface areas were close to the interpolated hexagon areas, and in all cases differed by less than 15%. Given that the expected facet area was interpolated from a single parameter vs. the 6 2D coordinates that are used for hexagon interpolation, we expected the area of the former would be more accurate. Hence, we applied a correction to the interpolated hexagon at each SP by scaling it such that its area equalled the expected facet area. This also adjusted the distance to the adjacent facet centres, and the average of all six distances was used as the local *D* of a given SP. Dividing the eye’s total surface area by the average expected facet area from all SPs indicated the total facet number of each compound eye.

### Corneal projection

By rotating the eye such that the NV at a given SP faced forwards, we could then select the corneal voxels that were closest the centres of its six adjacent facets determined from hexagon interpolation. The NV at each facet centre was then calculated, and the angle between the NV of the SP and each adjacent NV indicated the divergence angle between the corneal orientation of those facets. These six angles provide two measurements for each of the x-y-z axes of an individual facet (Land and Eckert 1985). The average of the six angles for a SP indicated the local corneal IO angle (*ΔΦ*), while the average of dividing *D* at each SP by the associated *ΔΦ* (in radians) indicated the local radius of curvature (*R*, Fig. 2B). Note that when calculated from corneal anatomy, *D* and *R* are typically measured first and the (corneal) IO angle is then obtained by calculating *ΔΦ*=*D*/*R* (Schwarz et al. 2011, Bergman and Rutowski 2016, Dyer et al. 2016), whereas we obtain *ΔΦ* and *D* first and calculated *R*=*D*/*ΔΦ*. The reason for this discrepancy is that we were not able to implement a satisfactory method for reliably measuring the local radius directly from the corneal surface voxels. Radius of curvature is a function of the 1^st^ and 2^nd^ derivatives of a surface whereas the NV is based only on the 1^st^ derivative, and calculating the latter appeared to be more numerically stable after the discretization of the eye into voxels.

The local eye convergence (*C*) was determined by calculating the 2D area enclosed by the hexagon of adjacent facet centres and dividing this by the solid angle subtended by the NVs projected from those centres. The corneal field of view (*FOV*) was calculated by dividing the eye’s total surface area by the average eye convergence of all SPs: 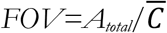, where *A_total_* is the total eye surface area.

Multiplying the corneal IO angle of each SP by *D* provided the local eye parameter (*P*): *P*=*ΔΦ*×*D*. Finally, the local ommatidial axis density (σ) projected into visual space was calculated for each SP by multiplying the eye convergence by the inverse of the interpolated facet area: *σ*=*C*/*A_facet_* where *A_facet_* is the facet surface area.

### Sensitivity calculation

The local optical sensitivity (*S*) of an ommatidia can be calculated as *S*=*A_facet_*×*Δϱ*^2^×(*k*×*L*/(1+*k*×*L*)) (Warrant and Nilsson 1998), where *A_facet_* is the facet surface area, *Δϱ* is acceptance angle of the ommatidia, *L* is the rhabdom length, and *k* is absorption coefficient of arthropod photoreceptors (0.0067 *μ*m^−1^). We could not measure local *Δϱ* or *L* directly in this study and approximated these by 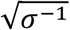 and retinal thickness respectively. While this substitution assumes that the acceptance angle is equal to the IO angle, it is often somewhat larger (Land 1997) and so our sensitivity calculation may underestimate absolute optical sensitivity.

### Determining fields of view

We generated a set of 10,242 world points (WP) equally spaced at approximately 1 ° intervals across the surface of a sphere that provided a common reference frame for world-centric calculations. The WPs within each bee’s FOV were determined with respect to the sphere intersects of the NVs from the SPs and BPs. Each SP was associated with a corneal IO angle, and we used nearest-neighbour interpolation to find an IO angle value for each BP from the SPs. We then tested whether each WP was within one IO angle range of the sphere intersection of any SP or BP, if so, they were included in our preliminary assignment to the set of WPs within the eye’s visual field (*FOV_left_*). The visual field was assumed to cover a single contagious region of the visual sphere, so any unassigned WPs that were completely enclosed were also assigned to *FOV_left_*. After these assignments, the fraction of the sphere covered by *FOV_left_* always exceed the calculated value for the corneal FOV by 5% – 10%. To resolve this discrepancy, we sequentially removed WPs from the periphery of *FOV_left_* until the angular area covered by both visual field representations was equal. *FOV_left_* was then mirrored to the opposite side of the visual sphere (and thus the frontal plane through the bee’s eyes, as they were aligned to face directly forwards), which represented the FOV of the bee’s right eye (*FOV_right_*). The binocular FOV was defined as the intersection of *FOV_left_* and *FOV_right_*, while the union of these sets of points define the complete FOV.

### Interpolation variables onto world points

To link the topology of variables between eye-centric and world-centric reference frames, we used IDW interpolation (with *p*=4) to interpolate each locally measured variable from the BPs and SPs to the WPs based on the angular distance between their sphere intersection points. The value of variables at the BPs were again found from nearest-neighbour interpolation from the values at SPs. The method of assigning values to the BPs forced the IDW interpolator to act as a nearest-neighbour interpolator at the periphery of the visual field, where it was effectively extrapolating from the SPs measurements. The variables interpolated to the WPs were: facet diameter, corneal IO angle, radius of curvature, thicknesses (lens, CC, and retinal), eye parameter, and ommatidial axis density.

### Validation of calculations

To validate the FOV calculation and corneal projection, we integrated the projected axis density across the WPs in the FOV of a given bee. This integral predicted the total number of ommatidial axes projected from the eye, which was compared to the total number of facets found from the area-based measurement. We considered the number of facets to be relatively accurate (in comparison to a study on honeybee corneal topography by Streinzer et al. (2013) we slightly overestimate the number of facets by less than 5%), while we knew that the choice of local radius influenced the NV calculation, which was the basis for calculating *ΔΦ* and *σ*, and must therefore influence the total axes number. If all calculations were completed using a small local radius (50 *μm*) substantially more axes were predicted than the number of facets, while fewer axes were predicted as the local radius was increased. However, this increase did not result in the axes number converging to the number of facets, as a large local radius (500 *μ*m) lead to the calculation of an insufficient axes number. To identify a suitable local radius for the calculations on each bee’s eye, we implemented a procedure that initially used a 100 *μ*m local radius for all NV dependent calculations, and then iteratively adjusted that parameter and repeated all calculations until the axes number was between 95% and 100% of the facet number. The required local radius was between 100 *μ*m and 200 *μ*m for most bees. The axes number was chosen to slightly underestimate the facet number to compensate for our expectation that the latter value slightly overestimated the true number of facets on an eye.

### Weighting means, distributions, and correlations

We compute mean values, relative distributions, and correlation coefficients for all local variables measured on each eye. The SPs at which all eye-centric variables are calculated are equally spaced across an eye while the facet density varies across it and simply using the values for each SP directly would misrepresent the calculation of these statistics. Therefore, we calculated weighted means, distributions, and correlations, where the weight used for each SP was the inverse of local facet surface area. This weighting represented the relative facet density of a SP and was analogous to performing statistics as if facet on the eye had been measured individually.

### Representing world-centric variables

Visual fields and the projected topology of any variable could be displayed either directly on a sphere (Fig. 2D) or as a 2D projection (Fig. 2E,F). In the latter case, we preferred to use a sinusoidal projection which preserves the representation of area on a sphere (Fig. 2F) but does not preserve the bearing between different points. Given the azimuth (*a*) and elevation (*e*) of a point on the sphere, the map coordinates (*x,y*) of a sinusoidal projection are calculated as: *x*=*a*×cos(*e*), and *y*=*e*.

We also calculated profiles of variable averages and integrals across bands of azimuth or elevation. To do this, we computed the azimuth and elevation of each WP, and then calculated the mean (or integral) for all WPs a bee viewed within a specific 10° range of either azimuth (for example, 0°<azimuth<10° | −90°<elevation<90°) or elevation (for example, 0°<el.<10° | −180°<az.<180°). Note that each azimuthal band had the same number of points, while the number of points in an elevation band decreased at the poles (el. ±90°). Averages were calculated as weighted means for the reasons given in the preceding section, with the exception being that WPs were weighted based on *σ*.

**Upon acceptance of this manuscript, we will upload the calculated data for each eye to the Datadryad repository and provide Ascension numbers to access these data. The Matlab scripts written to perform this analysis are available on request**.

### Comparison to the results of other studies

In our discussion, we compare the measurements from this study directly to the results of several others. We calculated ‘worst case’ errors when comparing our results to other studies on honeybees, as workers of this species have a highly consistent body size. ‘Worst case’ error was calculated as a percentage from our measurement that was furthest from the parameter reported in another study, plus one standard deviation in the opposite direction. For example, compared to the findings of Streinzer et al. (2013), both of our measurements for honeybee facet number (5440 and 5484) are larger than the reported values (5375±143), hence, we calculate the percentage error as 5484/(5375-143) =+4.8%.

### Limitations of our technique

We acknowledge that there are several limitations of our technique and these primarily relate to our inability to measure the internal dimensions of each ommatidia across the eye. To provide a calculation of sensitivity, it is necessary to determine the local rhabdom diameter and the focal length of the lens (Warrant and Nilsson 1998). Rhabdoms generally could not be clearly visualized in our imaged volumes (Figs. 1 & 2); their diameter was probably close to the voxel size (1.6 *μ*m), and limited contrast is seen between soft tissue structures. The focal length is an optical property of a lens and requires knowledge of its inner and outer radii, its thickness, and its refractive index to be calculate (Born and Wolf 1999). Although our measurements show lens thickness varies across each bee’s eye, suggesting that the focal length is likely to vary, the raw volumes did not have sufficient resolution to allow us to determine the outer and inner lens radii of individual facets on the eye (note that these are *not* equivalent to radius of curvature calculated in relation to IO angle). Improving the resolution and contrast with which the entire eye is imaged would allow the rhabdom diameter (and length) as well as the corneal lens structure of individual ommatidia to be measured and used in optical calculations.

Additionally, our descriptions of FOV and IO angles are at the corneal level and assume that the optical axes of ommatidia are perpendicular to the cornea. However, many studies have shown that the optical axis of ommatidia are often skewed from the lens normal (Stavenga 1979), and extreme differences of up to 40° have been reported in the ventral rim of honeybee eyes (Baumgärtner 1928). In such cases the visual field can be expanded at the cost of reduced resolution, and possibly sensitivity (the effective lens diameter is reduced and corneal reflections increase, although these may be offset by an increased acceptance angle). We could visualize the CC across parts of some compound eyes (Fig. 1B,C), but were unable to reliably segment them individually which would be required to incorporate their skewness into our analysis. Hence, we strictly use the terms *corneal* FOV and *corneal* IO angle; we believe these should provide a suitable basis for comparison between subjects that are likely to have similar patterns of CC offsets (such as between bees), although this assumption may not hold if comparisons are made between species with particularly different eye shapes (between the oval eyes of bees and the spherical eyes of butterflies for instance).

An additional limitation of this method is that we calculate an average IO angle at each sampling location on a compound eye, and do not decompose this into partial angles for the facet’s vertical and horizontal axes (Stavenga 1979). The resolution of many compound eyes is asymmetric, in that resolution in the vertical and horizontal directions differs (Land 1997). Sampling asymmetry occurs because either the hexagonal lattice of ommatidia is elongated or the eye’s radius of curvature varies directionally, and bee’s oval eyes are known to have higher vertical resolution than horizontal resolution in their frontal visual field (Seidl 1982, Spaethe and Chittka 2003). In this study we chose to average IO angles as we often found that their peripheral facets were packed irregularly and did not provide an obvious choice for facet axes, or that the facet axes were misaligned from the world axes (also noted by Seidl and Kaiser (1981)). Both factors complicate the calculation of partial IO angles across the entire visual field in a manner that would be analogous to the decomposed angles presented elsewhere, yet the average angle is directly comparable as this can be calculated from the vertical and horizontal angles reported in other studies.

Finally, we noticed that the relative accuracy of our method of calculating IO angles declined on the flattest compound eye areas. This was because the discretization of an imaged eye’s surface into voxels made areas with a large radius of curvatures indistinguishable from flat planes. We determined IO angles by firstly fitting polynomial surfaces to the voxels from small areas on the surface, and then calculating the angle between the NVs from points on these functions. If the function is fitted to a flat plane, then the NVs from all points across it will lie in parallel (indicating a misleading corneal IO angle of 0°). To prevent such inaccuracies, we implemented a heuristic that enlarged the local radius used to select the voxels used when fitting functions until it selected a curved set of points. Nonetheless, it is apparent that the largest relative errors for this method of calculating IO angle are likely to occur in the highest resolution areas of an eye (below 1° corneal IO angle), as these are also likely to be the flattest.

## Footnotes

1 Using a power function (*Y*=*bx^a^*) to relate a trait (*Y*) to a measure of body size (*x*) provides two parameters that describe allometry, the scaling exponent (*a*) and the initial growth index (*b*); Huxely & Teissier (1936).

2 With the relationships found in this study for ITW vs. EV, and for EV vs. facet number of Bombus terrestris workers (Table S1), we estimated the number of facets for bumblebees with the smallest and largest thorax widths reported in Fig. 4 of Spaethe & Chittka (2003), and also calculated the IO angles from the partial IO angles reported. The bee’s with ITW between 2.8 to 3.0 mm were estimated to have 4968 facets (from ITW 2.9 mm) and calculated to have an average full IO angle of 1.52°. For the four bees with 4.0 to 4.2 mm ITW, we estimate 5463 facets (from ITW 4.1 mm) and calculate an average full IO angle of 1.23°.

3 The projected area of a 3D object depends both on its shape and the direction of the observer as, for example, a hemisphere viewed along its midline appears as a circle with a projected area that is 50% of the true area. If the projected area of compound eyes in this study are calculated from orthographic projections along each corneas principle axis, the true surface area is consistently underestimated by 22%.

4 We calculated the ‘worst case’ error to be −4.4% for surface area, +4.8% for facet number, and +2.8% for maximum facet diameter.

5 Using the mean values for the eleven Bombus species reported in Table 1 of Streinzer and Spaethe (2014), we calculated the scaling exponent for both facet number and maximum facet diameter vs. √eye area. The scaling exponent for the maximum facet diameter vs. 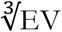 for the B. terrestris used in this study is 0.70, which is close to the scaling exponent we calculated for mean facet diameter (0.71). Note that the same exponent will be found whether √eye area or 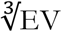 is used as the independent variable, although this choice influences the initial growth index.

6 Using the equation: *Angle*(°)=17.6−3.1×*ITW*(mm) from Spaethe and Chittka (2003). For our very small and large bumblebees (ITW 2.0 mm and 5.4 mm), this predicts visual angles of 11.5° and 0.7° respectively. These values appear unrepresentative of the data shown by Spaethe and Chittka (2003), suggesting that relationship between ITW and visual acuity may only be linear across the intermediate range of *Bombus* sizes tested in that study.

7 The influence of rhabdom length on sensitivity is as multiplier of *k*×*L*/(2.3+*k*×*L*), where *k* is absorption coefficient of arthropod photoreceptors (0.0067 *μ*m^−1^) and *L* is the rhabdom length in *μ*m (Warrant & Nilsson, 1998). We determined the ratio between the multiplier when calculated with the minimum and maximum retinal thickness on each eye.

## List of abbreviations

az.: Azimuth
CC: Crystalline cone
el.: Elevation
EV: Eye volume
FOV: Field of view
ITW: Inter-tegula width
IO: Inter-ommatidial
microCT: micro-computed tomography
NV: Normal vector

